# Synthetic mycobacterial molecular patterns partially complete Freund’s adjuvant

**DOI:** 10.1101/699165

**Authors:** Jean-Yves Dubé, Fiona McIntosh, Juan G. Zarruk, Samuel David, Jérôme Nigou, Marcel A. Behr

**Author notes:** Contact Info (Lead Contact) McGill University Health Centre 1001 boul. Décarie Glen Site Block E, Office #EM3.3212 Montréal, Québec, Canada H4A 3J1.

## Abstract

Complete Freund’s adjuvant (CFA) has historically been one of the most useful tools of immunologists. Essentially comprised of dead mycobacteria and mineral oil, we asked ourselves what is special about the mycobacterial part of this adjuvant, and could it be recapitulated synthetically? Here, we demonstrate the essentiality of *N*-glycolylated peptidoglycan plus trehalose dimycolate (both unique in mycobacteria) for the complete adjuvant effect using knockouts and chemical complementation. A combination of synthetic *N*-glycolyl muramyl dipeptide and minimal trehalose dimycolate motif GlcC14C18 was able to upregulate dendritic cell effectors, plus induce experimental autoimmunity qualitatively similar but quantitatively milder compared to CFA. This research outlines how to substitute CFA with a consistent, molecularly-defined adjuvant which may inform the design of immunotherapeutic agents and vaccines benefitting from cell-mediated immunity. We also anticipate using synthetic microbe-associated molecular patterns (MAMPs) to study mycobacterial immunity and immunopathogenesis.

## Introduction

Infection with *Mycobacterium tuberculosis*, or administration of bacille Calmette-Guérin (BCG), normally leads to cell-meditated immunity (CMI) to the corresponding bacterial antigens. The tuberculin skin test is positive when the immune response, a type IV hypersensitivity reaction or delayed-type hypersensitivity (DTH), occurs to tuberculin (a protein extract of *M. tuberculosis*). This “cutaneous sensitivity” was closely examined in the 1940s using complete Freund’s adjuvant (CFA, heat-killed *M. tuberculosis* in mineral oil plus surfactant) ^1, 2^. These studies provided the first direct evidence for the cellular nature of DTH by transfer from a CFA-immunized guinea pig to a naïve one only through the washed, heat-liable cellular fraction of peritoneal exudates ^1, 2^. It is now known that DTH is mediated specifically by antigen-sensitive T cells.

Today, CFA is a ‘gold standard’ adjuvant for eliciting CMI in research models of autoimmune disease. Notable is the experimental autoimmune encephalomyelitis (EAE) model of T-cell meditated destruction of myelin causing ascending paralysis, used most often to model multiple sclerosis ^3, 4^. CFA is not used in humans because of high reactogenicity ^5^. We have asked ourselves: what was the impetus for Jules Freund to develop his eponymous adjuvant with *M. tuberculosis*? For those conducting TB research, it is well appreciated that handling *M. tuberculosis* (a slow growing, clumping, fastidious, and lethally pathogenic organism) is a significant task *per se*. So, why did Freund choose to incorporate these bacteria?

Effort has been made to describe the microbe-associated molecular patterns (MAMPs) in mycobacteria that drive the adjuvant effect. Evidence has pointed to mycobacterial peptidoglycan (PGN), specifically down to the molecular motif muramyl dipeptide (MDP) ^6, 7^. Mycobacteria are distinct in that they produce the *N*-glycolyl MDP motif in their cell wall ^8–10^ and this PGN modification has been shown to be more potent compared to the common *N*-acetyl MDP motif possessed by other bacteria ^7, 11^. MDP is thought to be recognized through the host molecule NOD2 ^12^, and mutations in NOD2 predispose humans to increased risk of mycobacterial and inflammatory diseases ^13–16^. Alternatively, others have pointed to the mycobacterial cell wall lipid trehalose-6,6-dimycolate (TDM) alone or synergistically with purified peptidoglycan ^17^. TDM is recognized by the host with the C-type lectin Mincle in 18,19 concert with FcRγ and MCL. TDM has been demonstrated in animal models to alone be sufficient for granuloma formation and immunopathological responses ^18, 20^.

Both *N*-glycolyl MDP and TDM are MAMPs unique to mycobacteria. Their role in mycobacterial immune responses is supported by the literature and their host receptors are well-known. Additionally, *N*-glycolyl MDP is producible synthetically ^7, 21, 22^. Recently a minimal motif of TDM called GlcC14C18 was produced synthetically and shown to retain adjuvancy but with minimal toxicity ^23^. As a complex biologic, CFA is subjected to batch inconsistency. We hypothesized that it is possible to create an entirely synthetic CFA using a rational approach to identify essential MAMPs to replace the whole mycobacteria in the adjuvant. In this work, we establish the necessity for *N*-glycolylation of PGN, NOD2 and Mincle for full CFA adjuvancy. We also demonstrate that the mycobacteria in CFA can be replaced with synthetic *N*-glycolyl MDP and GlcC14C18 in combination to partially restore the adjuvant effect in both a murine model of antigen-specific T-cell immunity as well as the EAE model of autoimmune ascending paralysis.

## Results

### *N*-glycolylation of PGN by mycobacteria is required for complete adjuvancy

Previous work indicated mycobacteria required *N*-glycolylated PGN to elicit a maximal immune response during live infection ^7, 11^. To determine if CMI elicited from dead mycobacteria in the context of Freund’s adjuvant similarly required *N*-glycolylated PGN, we prepared in-house complete Freund’s adjuvant with heat-killed *M. tuberculosis* strain H37Rv, and H37Rv Δ*namH*. NamH is the enzyme responsible for *N*-glycolylation of PGN units as they are being synthesized in the cytoplasm, and the Δ*namH* mutant was previously shown to be devoid of *N*-glycolylation ^11^ (fig. 1A). We next immunized mice against ovalbumin (OVA, an exemplary antigen) with our CFAs or incomplete Freund’s adjuvant (IFA, lacks mycobacteria), and examined OVA-specific cytokine production by CD4+ T-cells from the draining lymph nodes seven days post-immunization (fig. 1B; gating in fig. S1A). Both mycobacteria and *namH* were required to generate the highest proportion of OVA-specific IFN-γ -producing CD4+ T cells (fig. 1C). OVA-specific IL-17A-producing cells required mycobacteria but did not significantly depend on *namH* (fig. 1D). Similar results were seen when examining total number of cytokine-producing CD4+ T cells in the lymph nodes (fig. S2A-B), and a closer analysis of the pooled data showed that *namH* contributes to about one third of mycobacteria-induced antigen-specific IFN-γ -producing Th cells (fig. S2C-D). The individual results of experiments are shown se tely in fig. S2E; in all four cases we saw the same trends with IFN-γ. These results are para consistent with the hypothesized role of mycobacteria being a key ingredient in CFA for eliciting CMI, and that mycobacterial PGN modification by NamH makes a more potent adjuvant.

**Figure 1.**
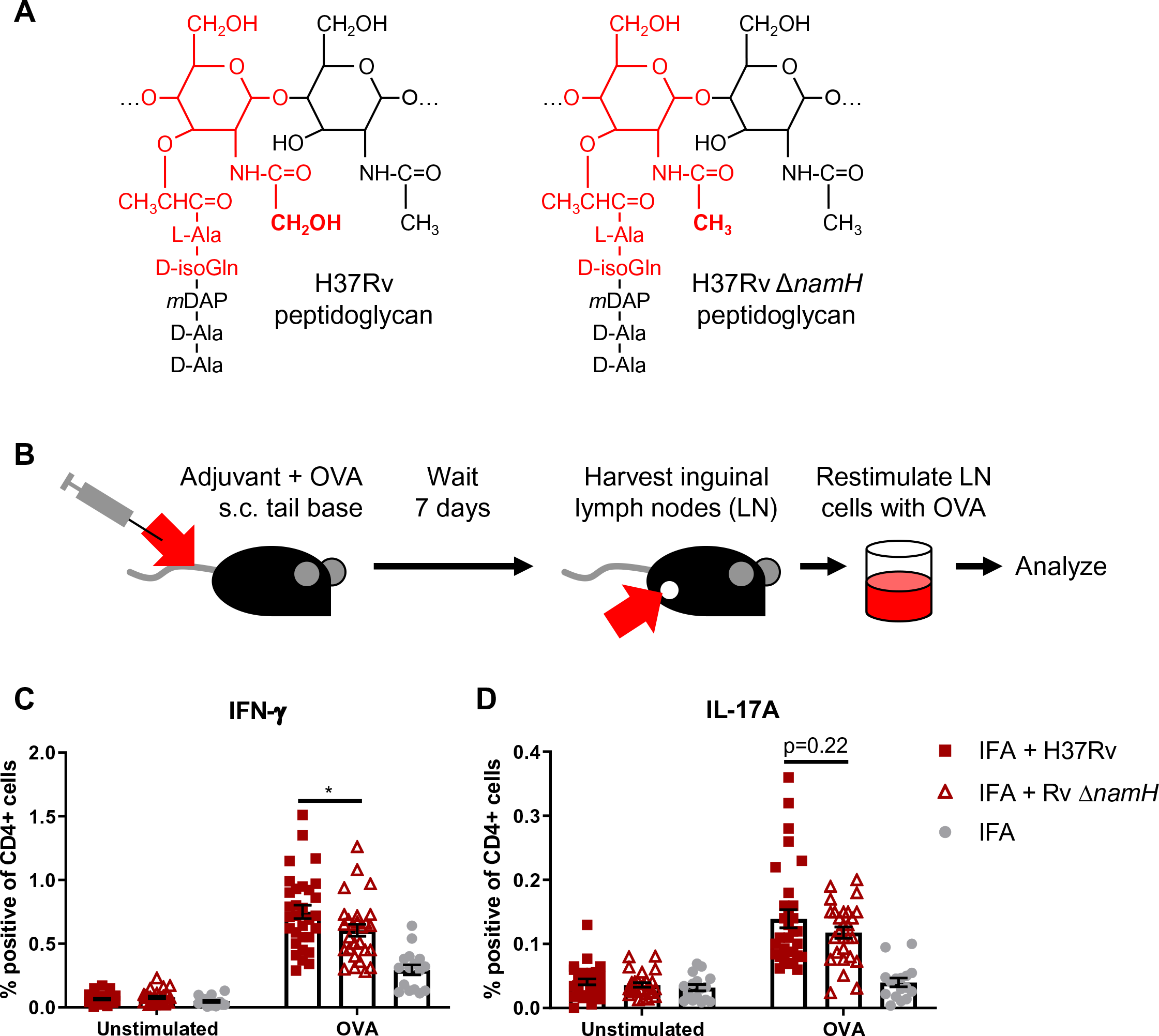
CFA-dependent cell-mediated immune responses as a function of mycobacterial Δ*namH*. **A**, PGN of wild-type H37Rv *M. tuberculosis* (left) and PGN of the (right). The MDP motif is drawn in red, and the site of *N*-glcolylation is in bold font. With NamH, *N*-glycolylation was shown on ∼70% of muramic acid residues, with *N*-acetylation on the remaining ∼30% ^8, 11^. **B**, immunization scheme (relevant to figs. 1, 2 and 4): mice were immunized with adjuvant emulsion containing OVA by s.c. injection at the base of the tail, and after seven days, inguinal (draining) lymph nodes were harvested. Lymph node cells were cultured *ex vivo* with or without OVA to examine the OVA-specific cytokine response by flow cytometry or ELISpot. **C-D**, Proportion of cytokine-producing CD4+CD8-lymph node cells of mice immunized against OVA with heat-killed *M. tuberculosis* strain H37Rv, H37Rv Δ*namH* or IFA alone, seven days prior. Shown are data pooled from four separate experiments with averages +/-SEM. p-values were calculated with two-tailed student’s *t*-tests. *p<0.05. For IFA Δ*namH*, and IFA alone, sample size N = 31, 27 and 16 mice, respectively. Each plott point represents the result obtained from an individual mouse.

### Host *Mincle* and *Nod2* are necessary for complete mycobacterial adjuvancy, but do not mediated the entire CFA effect

Recognition of mycobacteria during live infection requires the host molecule NOD2 ^24^. Because we hypothesize mycobacterial PGN plays a role in CFA adjuvancy, specifically the MDP motif where *N*-glycolylation occurs, we addressed whether the PGN/MDP sensor NOD2 is important for CFA adjuvancy in our OVA model. *Nod2*+/+ and *Nod2*-/-mice were immunized against OVA in the context of CFA or IFA of commercial provenance. A greater proportion of OVA-specific IFN-γ-producing CD4+ cells was generated in CFA-immunized mice with *Nod2* than without (fig. 2). An independently produced IFN-γ ELISpot of total lymph node cells A supported the same conclusion (fig. S3A), as did another independent experiment using flow cytometry (fig. S3C). In the absence of *Nod2*, the proportion of OVA-specific IL-17A-producing cells was also impaired in CFA-immunized mice (fig. 2B and fig. S3C). With IFA as adjuvant, IFN-γ and IL-17A responses were *Nod2*-independent as expected (fig. 2A-B and S3A-C). Similar trends were observed with total cell numbers (fig. S3B-C and S4A-B). Together, these results corroborate the role for NOD2 and accordingly mycobacterial PGN in the ability of CFA to elicit CMI.

**Figure 2.**
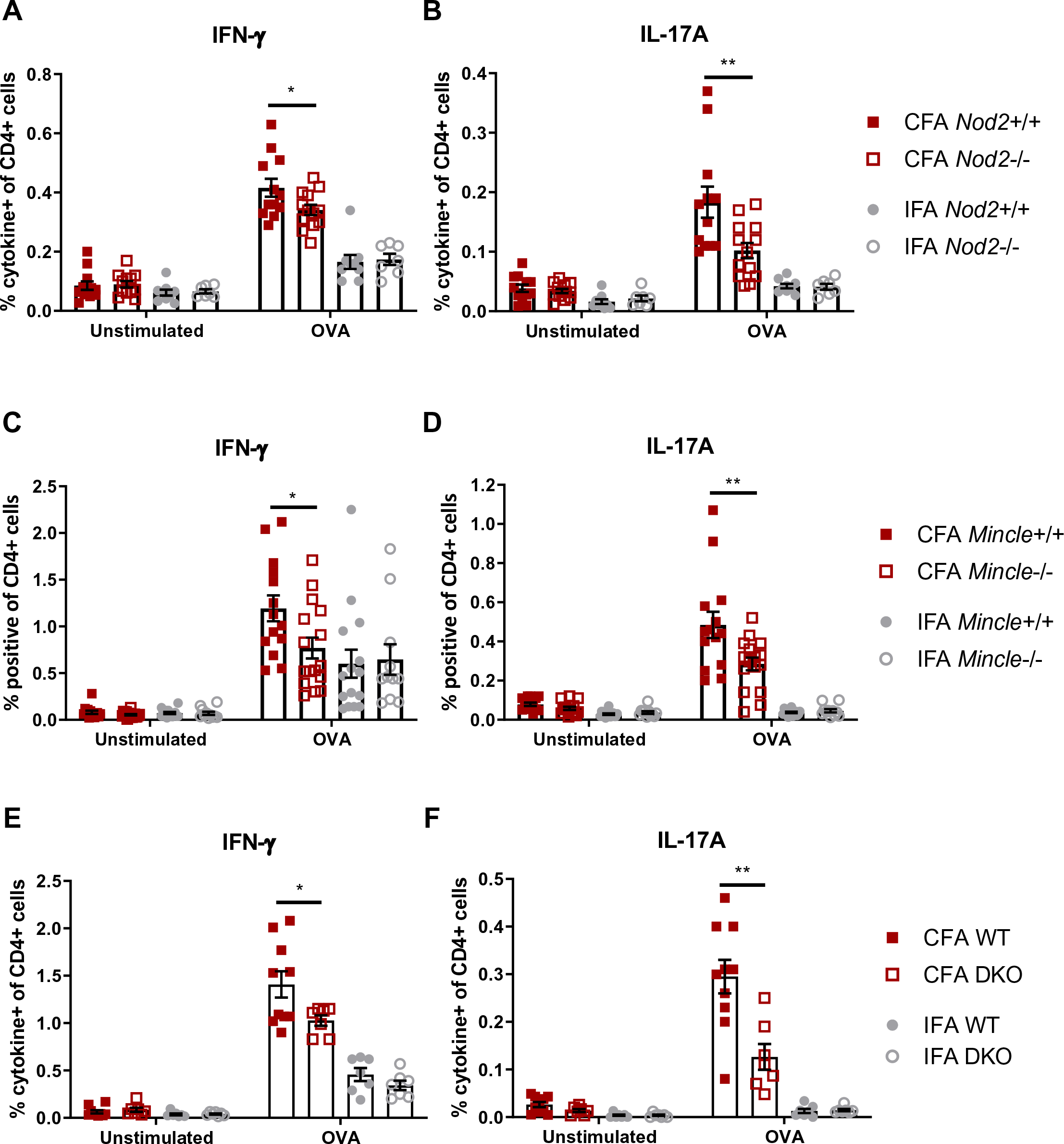
CFA-dependent cell-mediated immune responses as a function of host *Nod2* and *Mincle*. Proportion of cytokine-producing CD4+CD8-lymph node cells of mice immunized against OVA with CFA or IFA seven days prior. **A-B**, *Nod2*+/+ vs. *Nod2*-/-mice, data representative of two independent experiments with averages +/-SEM. p-values were calculated with two-tailed student’s *t*-tests. *p<0.05; **p<0.01. For CFA *Nod2*+/+, CFA *Nod2*-/-, IFA *Nod2*+/+ and IFA *Nod2*-/-, sample size N = 12, 13, 9 and 7 mice, respectively. **C**-**D**, *Mincle*+/+ vs. *Mincle*-/-mice, data pooled from two independent experiments with averages +/-SEM. p-values were calculated with two-tailed student’s *t*-tests. *p<0.05; **p<0.01. For CFA *Mincle*+/+, CFA *Mincle*-/-, IFA *Mincle*+/+ and IFA *Mincle*-/-, sample size N = 14, 16, 15 and 11 mice, respectively. **E**-**F**, WT (*Mincle*+/+*Nod2*+/+) vs. DKO (*Mincle*-/-*Nod2*-/-) mice with averages +/-SEM. p-values were calculated with two-tailed student’s *t*-tests. *p<0.05; **p<0.01. For CFA WT, CFA DKO, IFA WT and IFA DKO, sample size N = 10, 7, 7 and 7 mice, respectively.

The adjuvant effect of mycobacteria declined partially with *Nod2* knock-out, but not completely. Therefore, there is clearly other mycobacterial MAMP recognition besides that through NOD2 which is important for the remainder CFA adjuvancy. Others have indicated that Mincle-mediated recognition of TDM contributes to mycobacterial adjuvancy ^17^. We tested the dependence of our OVA-immunization model with *Mincle*-KO mice ^25^. *Mincle*+/+ and *Mincle*-/-mice were immunized against OVA in the presence of CFA and IFA. Generation of CFA-elicited OVA-specific IFN-γ- and IL-17A-producing CD4+ T cells was partially dependent on *Mincle* (fig. 2C-D), while I-immunized mice showed *Mincle*-independent responses, as FA expected (fig. 2C-D). Similar trends were again observed with total cell numbers but did not reach statistical significance (fig. S4C-D). Un-pooled data showed the same trends in both experiments performed (fig. S3D). These results implicate Mincle and thus its main mycobacterial ligand, TDM, in CFA adjuvancy.

Both *Nod2* and *Mincle* were required for CFA adjuvancy in our model, but neither were responsible for the entire adjuvant effect. We asked what the sum of the immune responses elicited by these PRRs is – if it appears these PRRs signal synergistically. We generated *Mincle*-*Nod2* double knock-out mice (DKO) and immunized them against OVA with CFA and IFA. The proportion of OVA-specific IFN-γ- and IL-17A-producing CD4+ T cells in CFA-immunized mice declined with DKO relative to WT mice, with no change in IFA-immunized mice (fig. 2E-F). However, the decrease when measuring IFN-γ did not appear obviously larger with DKO than what was observed with single-KO mice. L ewise, the decrease with IL-17A was intermediate, and similar trends were seen with total cell numbers (fig. S4E-F). The residual effect of CFA when both *Nod2* and *Mincle* are disrupted suggests that at least two mycobacterial MAMPs, *N*-glycolylated PGN as well as TDM, are essential, but insufficient, for the full adjuvant effect.

### Synthetic minimal structures of mycobacterial PGN and TDM work synergistically to stimulate dendritic cell effector functions

To explore the effect of Mincle and NOD2 signalling on antigen presenting cells (necessary to generate antigen-specific T-cell immunity), we stimulated bone-marrow derived dendritic cells (BMDCs) from WT, *Mincle*-/-, *Nod2*-/- and DKO mice with synthetic versions of the respective PRR ligands. Recently, a rationally determined minimal chemical structure of TDM called GlcC14C18 was shown to retain activity as a Mincle agonist without the ‘toxicity’ associated with purified TDM ^23^. Both synthetic *N*-glycolyl and *N*-acetyl MDP are commercially available. To elicit T-cell immunity, we would expect that mycobacterial MAMPs promote antigen presentation, co-stimulation and cytokine production by antigen presenting cells, the classical signals 1, 2 and 3, respectively.

After 24 hours of MAMP stimulation *in vitro*, we measured the proportion of cells in BMDC culture highly expressing MHC-II (fig. 3A-B). With WT cells, MDPs alone had little effect; GlcC14C18 alone increased the fraction of MHC-II^hi^ cells in WT BMDC cultures (fig. 3A). Combinations of GlcC14C18 with *N*-glycolyl MDP or *N*-acetyl MDP produced the largest responses. Knock-out of *Mincle* greatly reduced the effect of GlcC14C18 in *Mincle*-/- and DKO BMDCs, and knock-out of *Nod2* abrogated the effect of MDPs in *Nod2*-/- and DKO BMDCs. These results suggest these mycobacterial MAMPs can direct myeloid cells towards MHC-II^hi^ expression as expected for professional antigen presenting cells. Similar results were obtained at 48 hours of stimulation (fig. S5A).

**Figure 3.**
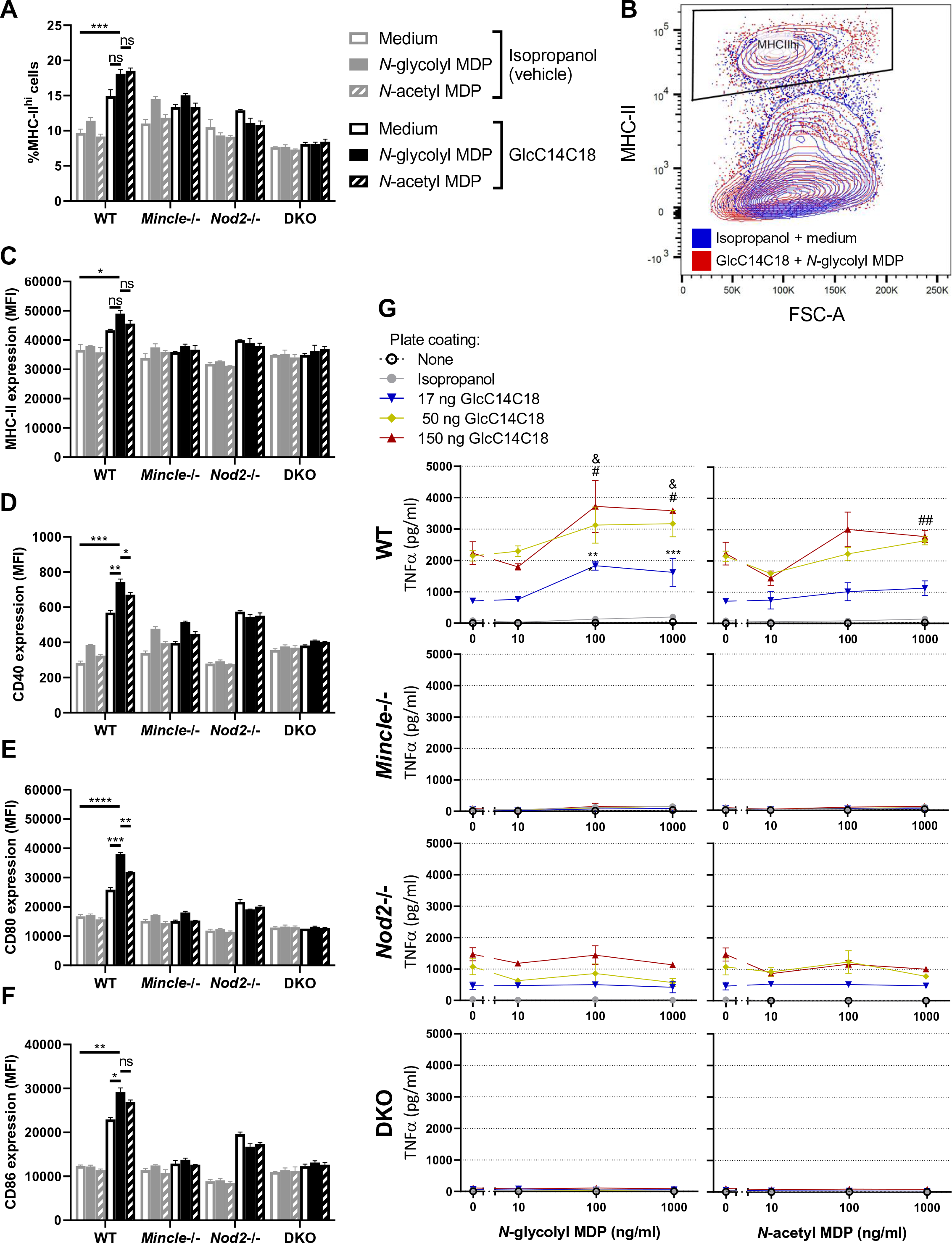
MHC-II expression, costimulatory molecule upregulation and TNF production and by BMDCs stimulated with GlcC14C18 and MDPs. **A**, percentage of cells expressing MHC-II at high levels (according to gate in panel B) after 24 hours of stimulation with the indicated MAMPs. **B**, gating of MHC-II^hi^ cells amongst live single CD11b+CD11c+ cells. **C-F**, median fluorescence intensity of **C**, MHC-II; **D**, CD40; **E**, CD80; **F**, CD86 on CD11b+CD11c+MHC-II^hi^ cells after 24 hours of stimulation with the indicated MAMPs (use legend of panel A). Shown are averages +/-SD of 3 individually stimulated and assayed cultures. To compare the combination of GlcC14C18+*N*-glycolyl MDP to unstimulated control, GlcC14C18 alone and GlcC14C18+*N*-acetyl MDP with WT cells, p-values were calculated using Dunnett’s T3 multiple comparisons test; p<0.05; **p<0.01; ***p<0.001; ****p<0.0001; ns, not significant, p>0.05. **G**, Supernatant TNF after 6-hour stimulation with GlcC14C18 and MDPs. Shown are averages +/-SD of 3-6 individually stimulated and assayed culture wells. To determine which MDP dose elicited significantly more TNF over background dose of GlcC14C18, p-values were calculated with Dunnett’s multiple comparisons test. The *, # and & symbols were used for the 17, 50 and 150 ng GlcC14C18 doses respectively, where *p<0.05; **p<0.01; ***p<0.001 (for clarity, non-significant p-values are not noted).

On MHC-II^hi^ cells, we looked at the level of expression of MHC-II, CD40, CD80 and CD86 by median fluorescence intensity (MFI) after 24 hours of stimulation *in vitro* (fig. S5F-I). With WT cells and for all markers examined, MDPs alone were unable to substantially enhance expression over the control (fig. 3C-F). GlcC1418 alone increased marker expression, but GlcC14C18 together with *N*-glycolyl MDP synergistically produced the greatest responses. *N*-acetyl MDP was generally less efficacious than *N*-glycolyl MDP. Knockouts behaved as expected: cells without *Mincle* responded poorly to GlcC14C18; cells without *Nod2* failed to respond to MDPs (fig. 3C-F). I addition to 24-hour stimulation, cells were also collected after 48 hours of stimulation, yielding similar results (fig. S5B-I).

BMDCs from WT mice produced TNF in a dose-dependent manner in response to GlcC14C18, but not to MDPs alone (fig. 3G). When combined with limiting or saturating doses of GlcC14C18, *N*-glycolyl MDP synergistically elevated TNF production in a dose-dependent manner. *N*-acetyl MDP only significantly elevated TNF production at higher doses of GlcC14C18 plus MDP and worked less efficaciously than *N*-glycolyl MDP. In the absence of functional *Mincle*, BMDCs did not respond to GlcC14C18 as expected. *Mincle*-/-BMDCs stimulated with GlcC14C18 and MDPs behaved like BMDCs exposed to MDP alone (little to no detectable TNF production). *Nod2*-/-BMDCs responded to GlcC14C18 but not to MDPs as expected. DKO BMDCs did not respond to either synthetic MAMP as expected. Overall, GlcC14C18 and *N*-glycolyl MDP combined, through *Mincle* and *Nod2* respectively, are sufficient to upregulate BMDC antigen presentation, co-stimulation and cytokine production.

### Synthetic mycobacterial NOD2 and Mincle ligands can complement IFA to increase antigen-specific T-cell responses

As NOD2 and Mincle ligands worked synergistically in BMDCs *in vitro* to elicit effector functions in these antigen presenting cells, we hypothesized that IFA can be complemented with NOD2 and/or Mincle ligands to recapitulate at least a portion of the adjuvant effect of whole mycobacteria. Using minimal synthetic MAMPs for PGN (i.e. *N*-glycolyl MDP) and TDM (i.e. GlcC14C18), we complemented IFA to test if CMI could be achieved with a completely synthetic adjuvant containing these essential mycobacterial MAMPs. First, using 10 µg GlcC14C18 per mouse with varying doses of *N*-glycolyl MDP, we observed a dose dependent increase in the proportion and numbers of OVA-specific cytokine-producing CD4+ cells which was significantly greater than IFA alone at the highest dose (fig. 4A). Notably, previous attempts complementing IFA with only *N*-glycolyl MDP did not induce significant numbers of IFN-γ producing cells (fig. S6A-B).

**Figure 4.**
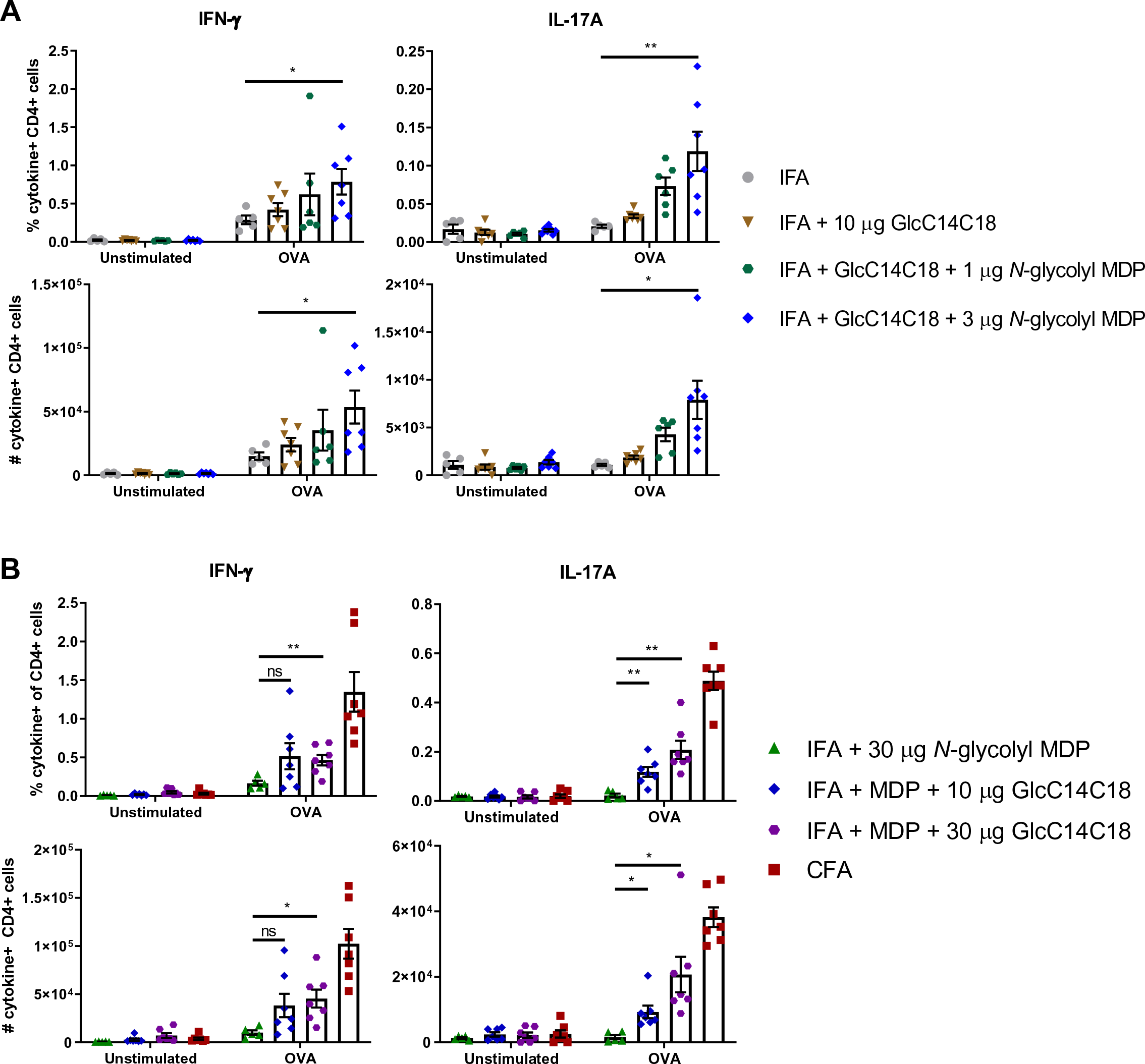
Complementation of IFA with synthetic mycobacterial MAMPs. Proportion of cytokine-producing CD4+CD8-lymph node cells of WT mice immunized against OVA seven days prior. **A**, *N*-glycolyl MDP dose-dependent response with 10 µg GlcC14C18: IFA (N=4), IFA + 10 µg GlcC14C18 (N=7), IFA + 10 µg GlcC14C18 + 1 µg *N*-glycolyl MDP (N=6), and IFA + 10 µg GlcC14C18 + 3 µg *N*-glycolyl MDP (N=7). Shown are data from individual mice, with averages +/-SEM. p-values were calculated with Welch’s two-tailed student’s *t*-tests. *p<0.05; **p<0.01. **B**, GlcC14C18 dose-dependent response with 30 µg *N*-glycolyl MDP: IFA + 30 µg *N*-glycolyl MDP (N=6), IFA + 30 µg *N*-glycolyl MDP + 10 µg GlcC14C18 (N=7), IFA + 30 µg *N*-glycolyl MDP + 30 µg GlcC14C18 (N=7), and CFA (N=7). Shown are data from individual mice, with averages +/-SEM. In comparing IFA + 30 µg *N*-glycolyl MDP to IFA + *N*-glycolyl MDP + 10 or 30 µg GlcC14C18, p-values were calculated using Dunnett’s T3 multiple comparisons test. *p<0.05; **p<0.01; ns, not significant, p>0.05.

Next, using 30 µg of *N*-glycolyl MDP per mouse and varying the dose of GlcC14C18, we similarly observed a dose dependent increase in the immune response; however this response did not surpass more than half of the level elicited with CFA by IFN-γ and IL-17A (fig. 4B). An IFN-γ ELISpot performed in parallel provided a similar conclusion (fig. S6C-D). Together, these results show that mycobacterial adjuvancy in the context of CFA can be partially phenocopied synthetically with two mycobacterial MAMPs: namely *N*-glycolyl MDP and GlcC14C18.

### Synthetic mycobacterial MAMPs increase DC numbers and effectors in lymph nodes

GlcC14C18 and *N*-glycolyl MDP together were sufficient to elicit antigen-specific T-cell responses. We next looked to see if this MAMP combination was associated with augmentation of DC effector functions *in vivo* as we had seen *in vitro* with BMDCs. Gating for DC subsets as well as other leukocytes in the lymph nodes is depicted in fig. S1B (guided by ^26^). We looked in the lymph nodes at 4- and 7-days post-immunization. Both CD11b+ and CD11b-conventional DC (cDC) numbers were increased in the presence of IFA + 10 μg GlcC14C18 + 30 μg *N*-glycolyl MDP compared to IFA alone within 7 days, to levels similar to CFA (fig 5A-B). Trends suggest that early accumulation of cDCs could be driven by MDP alone, but GlcC14C18 together with *N*-glycolyl MDP were required to sustain numbers to day 7.

By MFI, expression levels of MHC-I and -II on both CD11b+ and CD11b-cDCs were not largely altered by MDP nor GlcC14C18 (fig. 5A-B). On CD11b+ cDCs, CD40 expression was elevated the highest and significantly with IFA + GlcC14C18 MDP at both 4 and 7 days. CD80 and CD86 expression were also highest with IFA + GlcC14C18 MDP, with statistical significance at day 4 but not day 7 when expression was possibly waning. On CD11b-cDCs, CD80 and CD86 expression behaved similarly to that on CD11b+ cDCs. CD40 expression of CD11b-cDCs was not statistically significantly greater with IFA + GlcC14C18 MDP compared to IFA alone at both timepoints. Notably, IFA + GlcC14C18 MDP produced equal or greater costimulatory molecule expression compared to CFA in all cases.

**Figure 5.**
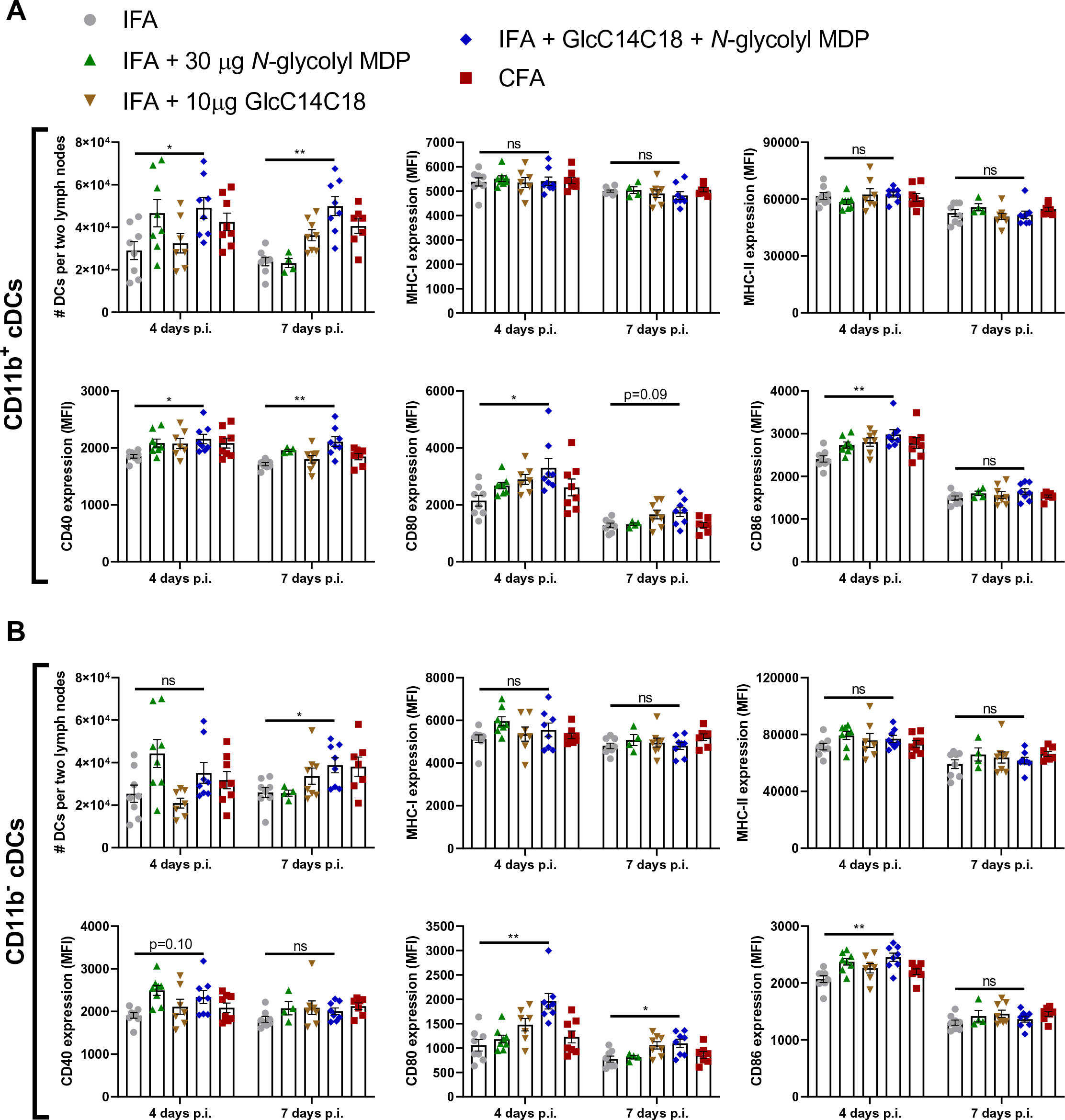
Dynamics of cDCs in lymph nodes after immunization with synthetic mycobacterial MAMPs. **A**, CD11b+ cDC quantities and median fluorescence intensity of MHC-I, MHC-II, CD40, CD80 and CD86 in mice immunized with the indicated adjuvants after 4 or 7 days. **B**, CD11b-cDC quantities and median fluorescence intensity of MHC-I, MHC-II, CD40, CD80 and CD86 in mice immunized with the indicated adjuvants after 4 or 7 days. For all adjuvant groups, N=8 mice (each plotted point represents data from an individual mouse). Shown are averages +/-SEM. To determine if IFA + GlcC14C18 + *N*-glycolyl MDP altered cDC parameters compared to IFA alone, p-values were calculated using Dunnett’s T3 multiple comparisons test correcting for multiple timepoints and cDC type (CD11b+/-). *p<0.05; **p<0.01; ns, not significant, p>0.05.

In addition to increased numbers of cDCs in the lymph nodes of mice immunized with GlcC14C18 + *N*-glycolyl MDP, we observed that other leukocytes were substantially elevated in numbers after immunization with this adjuvant (fig. S7A-F). B cells, CD4+ and CD8+ T cells, plasmacytoid DCs, monocytes and PMNs were all increased by 7 days post-immunization relative to IFA, to levels similar to or greater than CFA. By day 7, while MDP alone was insufficient to elevate cell numbers over the IFA background, MDP in combination with GlcC14C18 enhanced numbers beyond the levels attained with GlcC14C18 alone. This provides evidence for synergy between these MAMPs *in vivo*.

### Synthetic mycobacterial MAMPs can induce EAE similar to CFA

One of the common uses of CFA is in animal models of autoantigen-specific autoimmunity. To determine if our synthetic formulation can phenocopy the effect of whole mycobacteria to produce a more complex biological outcome such as autoimmunity (and thereby provide a further measure of the ‘completeness’ of the synthetic formula), we tested IFA + GlcC14C18 + *N*-glycolyl MDP against CFA in ability to induce relapsing-remitting EAE (RR-EAE). Briefly, mice were randomly immunized against myelin oligodendrocyte glycoprotein (MOG) synthetic peptide with CFA or IFA + 10 μg GlcC14C18 + 30 μg *N*-glycolyl MDP, and onset of RR-EAE was determined by clinically scoring ascending paralysis daily in a blinded manner (fig. 6A). Both CFA and IFA + GlcC14C18 + MDP produced RR-EAE that was indistinguishable except quantitatively: over the course of the experiment, the average EAE score was lower with the synthetic adjuvant compared to the CFA control (fig. 6B), with a cumulative score suggesting about half the disease burden (58% of CFA cumulative score) (fig. 6C). Expectedly, lower EAE scores also corresponded with less weight-loss (fig. S8A). Of note, there were mice in the synthetic adjuvant group that reached the same scores as in the CFA group, but fewer (fig. S8B). Therefore, GlcC14C18 and *N*-glycolyl MDP were sufficient to recapitulate the adjuvant effect of the whole mycobacterial cell in EAE, albeit quantitatively less at the tested doses of MAMPs. Additionally, we had attempted RR-EAE with IFA + TDM + MDP previously, which produced a far less compelling phenocopy of CFA (fig. S8D-G). The IFA + GlcC14C18 + *N*-glycolyl MDP adjuvant has the advantage of being completely synthetic and more efficacious than the TDM-containing adjuvant.

**Figure 6.**
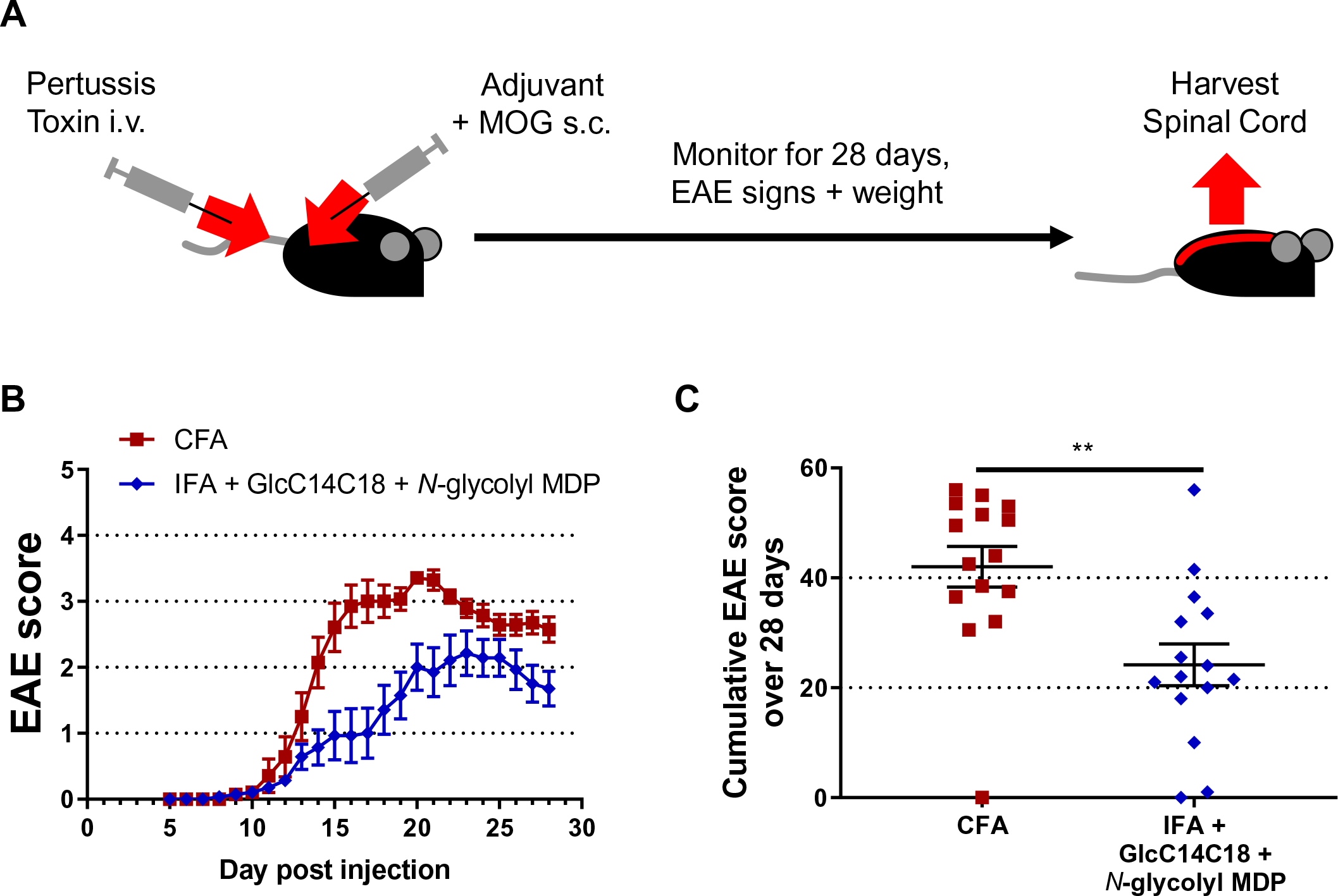
RR-EAE induced by IFA+GlcC14C18+MDP. **A**, experimental timeline. **B**, average EAE score +/-SEM over time of mice induced with CFA (N=15) or IFA + 10 µg GlcC14C18 + 30 µg *N*-glycolyl MDP (N=15). Mice were euthanized on day 28 post injection. **C**, Cumulative EAE score, obtained by adding the EAE score of each mouse over each of the 28 days. Lines represent averages +/-SEM. p-value was calculated with two-tailed student’s *t*-test. **p = 0.0022.

To verify if there were overt qualitative histopathological differences in EAE between CFA and the synthetic adjuvant, we examined spinal cords from three mice from each adjuvant group, having EAE scores 2, 3 and 3.5 at harvest (disease profile of these mice in fig. S8C). Both cellular infiltrates and demyelination of the white matter looked equivalent between adjuvants upon visual inspection (fig. 7A). Mice with different EAE scores had correspondingly different areas of diseased tissue of the spinal cord (fig. 7B), but in comparing adjuvants there was no detectable difference in the area of diseased spinal cord tissue (with statistical power to detect as low as +/-15% CFA levels) (fig. 7C). The main determining variable was EAE score, not adjuvant. Overall, we observed qualitatively similar autoimmune pathology using IFA + GlcC14C18 + *N*-glycolyl MDP compared to the gold standard CFA.

**Figure 7.**
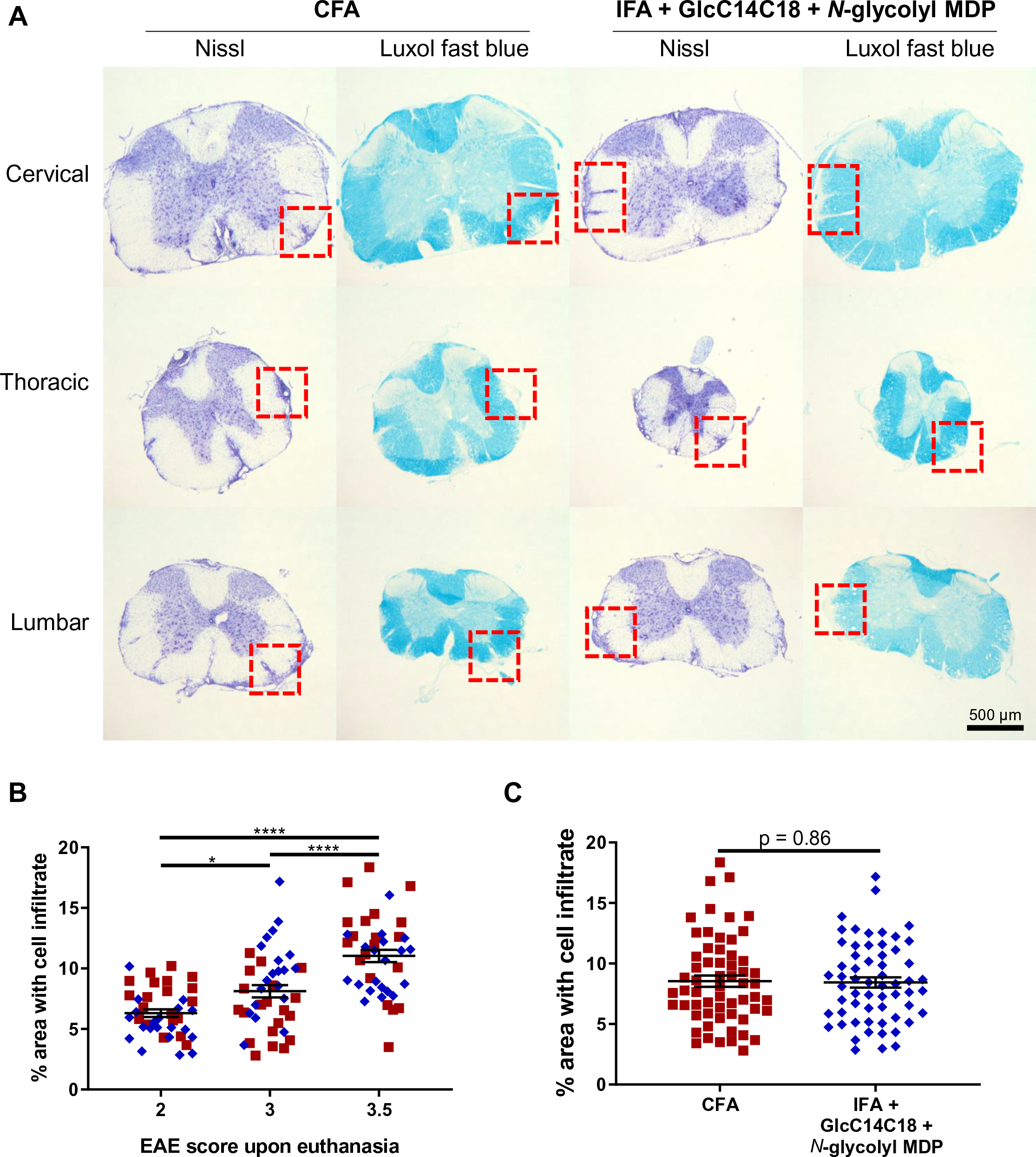
Spinal cord pathology in RR-EAE mice induced by IFA+GlcC14C18+MDP. **A**, Nissl and Luxol fast blue (LFB) stains of spinal cord sections from RR-EAE-induced mice at day 28 post injection, having an EAE score of 3.5 upon euthanasia. Red boxes highlight cellular infiltration and spatially associated demyelination of the white matter seen by Nissl and LFB staining, respectively. **B**, Quantitative spinal cord pathology per EAE score upon euthanasia. Statistical significance was determined by Tukey’s multiple comparisons test. *p<0.05 and ****p<0.0001. N=40 for each group (20 from CFA and 20 from synthetic adjuvant of equivalent EAE scores). Lines indicate averages +/-SEM **C**, Quantitative spinal cord pathology per adjuvant. Statistical significance was tested with two-tailed unpaired Welch’s *t*-test; a power calculation for given variances and N=60 per group indicated an ability to discern +/-15% difference vs. CFA control. Lines indicate averages +/-SEM.

## Discussion

The ability of an adjuvant to elicit CMI can depend on ligand-receptor interactions, specifically MAMP-PRR interactions. It is well appreciated since the hypothesis of Charles Janeway Jr. that the host decision to mount an adaptive immune response requires genetically inborn sensors to detect the presence of microbial products, or microbes by association, and that this is the foundation of classical adjuvants including CFA ^27^. These interactions can be thought of as an “arms race” between host and microbe, originating through antagonistic evolution in the case of immune evasion ^28^. Additionally, we can imagine the case where a microbe might evolve to increase a specific PRR interaction that favours a specific active immunological environment necessary for its lifecycle. When mycobacteria interact with their host, either in the form of *M. tuberculosis* infection, BCG vaccination or immunization using CFA, CMI normally occurs. The immune response to mycobacteria has been attributed to its unique cell wall, especially the MDP motif of PGN ^6, 7, 11, 24^ and TDM ^17, 19^. We have shown in the context of both CFA-induced immunization and autoimmunity with wholly synthetic *N*-glycolyl MDP and TDM (GlcC14C18) that these MAMP-PRR pathways together contribute partially but significantly to the mycobacterial adjuvant effect.

There are other mycobacterial MAMPs previously identified (and likely more yet unidentified), plus a greater number of PRRs linked to these microbial products. However, most are not yet producible synthetically. Purified mannose-capped lipoarabinomannan (ManLAM) was recently shown to elicit EAE in mice, dependent on the host C-type lectin Dectin-2 (Clec4n), at a dose of 500 µg per animal ^29^. Of note, we were not able yet to produce synthetic mimics of ManLAM that induce Dectin-2 signaling ^30^. Similarly, purified TDM has been used to elicit EAE at 500 µg per mouse, mostly dependent on MCL (Clec4d) and partially on Mincle (Clec4e) ^19^. These findings support a role for mycobacterial MAMPs driving the CFA effect, but require purification of biologically-sourced MAMPs in high doses. When in combination, our synthetically pure MAMPs have shown clear activity *in vivo* beginning at 1-10 µg. Of note, with CFA we inject 50 µg of mycobacterial mass into mice.

TLR-2 has been documented to interact with multiple mycobacterial lipids ^31^. The literature is inconsistent on whether or not TLR-2 promotes CFA-induced EAE ^32–35^. Mycobacterial DNA was recently shown to signal through the cGAS/STING pathway after phagosomal disruption via ESX-1 ^36, 37^. In the context of Freund’s adjuvant, it is not clear if heat-killed mycobacteria could have cytosolic access to activate cGAS/STING. The IFA fraction perhaps could deliver MAMPs across membranes. TLR-9 also senses DNA, and appears to play a role during *M. tuberculosis* infection in mice ^38^, and in humans ^39^. TLR-9 was also shown to be necessary for full induction of EAE ^32, 34^. Although DNA is not unique to mycobacteria like TDM and *N*-glycolylated PGN, synthetic DNA is available and pursuing the role of DNA in mycobacterial adjuvancy interests us, perhaps as a third synthetic MAMP to work with MDP and GlcC14C18.

Mycobacteria are not completely unique in possessing MAMPs that elicit CMI. As an example, the tuberculosis vaccine candidate M72/AS01E utilizes monophosphoryl lipid A (MPLA), a TLR-4 agonist based on lipopolysaccharide (LPS), to elicit CMI. LPS is made in Gram-negative bacteria, not mycobacteria. We concede that many microbes likely contain MAMPs able to elicit CMI; here we have concerned ourselves primarily with the history of mycobacterial adjuvancy, CMI and Jules Freund. It is conceivable that a combination of MAMPs from different bacteria could elicit ‘unnatural’ immunity that may be beneficial to control certain infectious agents that otherwise evade natural immune responses.

We were able to show that *N*-glycolylation of *M. tuberculosis* PGN altered the adjuvancy of the mycobacterial cell in the context of CFA. This is consistent with our previous work on the increased potency of mycobacterial PGN in other contexts ^7, 11^. Interestingly, the benefit of *N*-glycolylation of PGN seemed primarily skewed toward IFN-γ-production rather than IL-17A, while *Nod2*-KO showed a greater effect with IL-17A. This may indicate that mycobacteria retain *N*-glycolylation of PGN specifically to shift immune responses in a Th1 bias and should be further investigated during live infection. *Nod2* was necessary for full CFA adjuvancy by measures of IFN-γ and IL-17A, supporting the role for mycobacterial PGN. We were somewhat surprised that MD Pemulsified by itself with IFA and antigen was insufficient to recapitulate the adjuvant effect of CFA. In a 1975 paper, *N*-acetyl MDP appeared sufficient in a guinea pig model to elicit DTH using OVA as antigen ^6^. In 2009 using a very similar model to our current work, *N*-glycolyl MDP alone was sufficient to increase ELISpot IFN-γ to near CFA levels, while *N*-acetyl MDP failed ^7^. One explanation is the possibility of contaminating MAMPs in certain preparations of OVA. OVA is known to often come with a dose of endotoxin which affects the immunological outcome of experiments ^40^. In the current investigations we have used endotoxin-free/ultra-pure OVA. Much literature of *in vitro* experiments shows MDP requires another MAMP to produce measurable outputs like cytokine ^7, 41–43^, although in few cases MDP was shown to elicit cellular responses by itself ^41, 42^. Responses measured from MDP alone may be unrepresentative of a cell’s maximum capacity.

Other groups have supported a role for TDM in mycobacterial adjuvancy ^17, 19^. TDM together with purified PGN was shown to synergistically promote IL-17 production from OVA-specific (OT-II) CD4+ T cells adoptively transferred into congenic mice ^17^. This and other work directed us to test Mincle-dependence of CFA and MDP synergy with the minimal TDM motif GlcC14C18. *Mincle*-/-mice allowed us to infer that CFA adjuvancy required TDM for maximal induction of IFN-γ- and IL-17A-producing Th cells. TDM is thought to be the main Mincle ligand in mycobaria; however, there are other ligands. Purified from H37Rv, trehalose-6,6’-monomycolate, glucose monomycolate, diacyl-trehalose and triacyl-trehalose were shown to stimulate mouse and human Mincle (plus glycerol monomycolate stimulates human Mincle only)^23^. It is possible that the phenotype of reduced immunity was because the sensing of these other MAMPs was decreased. Nevertheless, Mincle signalling was essential for full CFA adjuvancy.

Interestingly, simultaneous knock-out of both *Mincle* and *Nod2* resulted in incomplete abrogation of CFA adjuvancy. DKO mice more closely resembled single-KO from separate experiments, compared to controls, than IFA. This indicates two things: 1) that much of the adjuvancy of mycobacteria is not mediated through Mincle and NOD2, and therefore other mycobacterial MAMPs like those mentioned above may play a similarly important role as PGN and TDM; 2) that Mincle and NOD2 signalling might work synergistically, where in the absence of Mincle, NOD2 sensing of PGN is blunted as we have seen clearly *in vitro* with GlcC14C18 and *N*-glycolyl MDP, and therefore DKO resembles single knock-out of either receptor.

We demonstrated that GlcC14C18 plus *N*-glycolyl MDP synergistically were sufficient to promote dendritic cell effector functions *in vitro*. The GlcC14C18 plus *N*-glycolyl MDP adjuvant was also the most efficacious adjuvant to drive costimulatory molecule upregulation on cDCs *in vivo*, more than either MAMP alone. While either of these MAMPs by themselves failed to come close to the whole mycobacteria of CFA in our OVA model of CMI, together we showed they could produce about half the Th cell IFN-γ /IL-17A response and EAE disease burden as CFA. Resultingly, we are interested in further understanding the implications for MAMP signalling (which MAMPs synergize together; what redundancy exists; do different combinations alter the Th type of CMI?). As the DKO data suggest, other mycobacterial MAMPs may be necessary to fully recapitulate the adjuvant effect of whole mycobacteria. An optimal dose of the two MAMPs may not have been reached, although additional dosing experiments (not shown) failed to find a clearly superior formulation than those presented here.

We chose to proceed primarily with the 10 µg GlcC14C18 + 30 µg *N*-glycolyl MDP doses because these quantities were sufficient for some biological activity on their own and/or in combination (fig. 4A-B, S6D and not shown). Detailed subset data also showed other biological synergy at the 10 µg / 30 µg dose (fig. S7). Mycobacteria are an estimated 30% PGN by mass, of which the muramyl dipeptide motif comprises about half, or 15% total mycobacterial mass ^44^. Therefore, when injecting 50 ug of mycobacteria as CFA, we inject about 8 µg of MDP. The 10 µg / 30 µg dose did not quantitatively phenocopy the level of CMI from CFA in the OVA model, but we did not want to achieve this by using grossly unrepresentative quantities of MAMPs. Considering the DKO data showing *Mincle* and *Nod2* stimulation do not drive all of the CFA effect, we proceeded with tempered expectations for EAE. Nonetheless, we have demonstrated that synthetic TDM and *N*-glycolyl MDP signalling, unique to mycobacteria, enhance the adjuvancy of CFA. This partly corroborates the natural history of CFA in why it is a uniquely potent adjuvant. Remaining MAMPs, not necessarily unique to mycobacteria, might fill the difference.

With the wholly synthetic formulation of GlcC14C18 *N*-glycolyl MDP, we were able to induce RR-EAE in mice that was qualitatively indistinguishable from that induced with CFA, but milder overall (lower EAE scores on average). We selected spinal cords from mice having matched EAE scores between CFA and synthetic adjuvant groups; however, extrapolating the histopathological data to all the mice in our experiment, we would expect that the synthetic adjuvant produced less pathology in the spinal cord on average since EAE scores were lower on average. The EAE result correlated with the incomplete complementation in the OVA model. EAE that failed with TDM plus MDP earlier was also less potent in the OVA model than the synthetic formulation (not shown), further indicating correlation between these readouts. CFA often needs to be ‘titrated’ to account for batch-dependent efficacy in our experience (not shown), and CFA with higher concentrations of mycobacteria (i.e. 4 mg/ml) is also used when a more severe, chronic progressive EAE is necessary for studies ^45^. These facts point to dosing as a means of controlling the degree of EAE, which may work with the synthetic adjuvant. Other experimental uses of CFA include the collagen-induced arthritis (CIA) model and for producing specific antibodies. CIA takes longer and has a greater role for humoral immunity than EAE which is T-cell driven, (our interest was in CMI for this study) ^46^. Wherever mycobacterial adjuvancy is useful, we suspect a synthetic adjuvant could be beneficial. Indeed, intravesical injection of BCG is used to treat bladder cancer (i.e. BCG can provide MAMP-driven protection, not simply antigen-specific protection). Furthermore, BCG administration to millions of babies each year can protect not just against childhood tuberculosis, but non-specifically against other diseases ^47^.

Our work demonstrates that mycobacterial NOD2 and Mincle ligands contribute to the adjuvant effect of the mycobacterial cell, and that a necessarily combinatorial formulation of synthetic MAMPs, *N*-glycolyl MDP and GlcC14C18 can partly recapitulate the adjuvant effect of whole mycobacteria to induce T-cell immunity and EAE. In addition to demonstrating the first entirely synthetic multi-MAMP mycobacterial adjuvant, we have outlined an approach to investigate the contribution of other MAMPs to adjuvant design. Moreover, the synthetic approach may be useful to probe mycobacterial immunity and immunopathogenesis.

## Methods

### Mice

C57BL/6J (‘wild-type’) as well as *Nod2*-/-mice were obtained from Jackson laboratories and were bred or used immediately for experiments. *Mincle*-/- mice breeders were provided courtesy of the laboratory of Christine Wells ^25^. Mice were genotyped to confirm absence of the *Dock2* mutation recently reported in *Nod2*-/- mice ^48^. We crossed *Nod2*-/- and *Mincle*-/- mice to obtain double knock-out mice. All experiments used mice from 6 to 20 weeks of age. All protocols involving mice followed the guidelines of the Canadian Council on Animal Care (CCAC) and were approved by the ethics committees of the RI-MUHC.

### Preparation of adjuvants and ovalbumin immunization

Complete and incomplete Freund’s adjuvant were purchased from Sigma or Invivogen. In some cases, IFA was made by the experimenter from purified mineral oil and mannide monooleate (Sigma). Adjuvants were prepared on the day of immunization by emulsifying CFA (1 mg/ml *M. tuberculosis*) or IFA with a PBS solution containing 1 mg/ml ovalbumin (Endofit brand, invivogen) in a 1:1 ratio. Emulsification was accomplished using all-plastic syringes and repeated passage through an 18-G blunt-end needle. Where IFA was complemented with mycobacterial components: MDP (Invivogen) was diluted in the PBS fraction of the adjuvant before emulsification; TDM (Sigma) or GlcC14C18 ^23^ were dissolved in the IFA fraction before emulsification as described previously ^17^; heat-killed mycobacteria in saline were diluted in PBS and added to the PBS fraction of the adjuvant before emulsification for mice to receive an equivalent of 10^8^ CFU. To make heat-killed mycobacteria, cultures were grown to equivalent 600-nm absorbance at mid-log phase, were pelleted and washed three times with saline, then heat-killed at 100°C for 30 minutes and frozen at −80°C until use. Mice were injected subcutaneously with 100 μl adjuvant-antigen emulsions in the tail 1-2 cm from the body, towards the body, for sufficient and consistent drainage to the inguinal lymph nodes. Four or seven days after injection mice were euthanized and organs were harvested for analysis.

### ELISpot and flow cytometry of lymph node cells

Lymph nodes extracted from immunized mice were gently crushed over 70-μm cell strainers to release the cells therein. Cell concentrations were determined by counting with a haemocytometer or with a BD Accuri flow cytometer. Lymph node cells (LNCs) were washed in culture medium, counted and cultured in 100 μl RPMI + 10%FBS (R10) at 250,000 or g OVA for ∼40 hours at 37°C 5% CO_2_ before developing the ELISpot plates. For flow cytometry analysis of OVA-specific cytokine production, washed LNCs were cultured at 6 million cells per ml in 200 μl R10 with or without 200 μg OVA for ∼40 hours at 37°C 5% CO_2_, Brefeldin A (GolgiPlug^TM^, BD) was added for an additional 5 hours, and then cells were stained and fixed (BD fixation/permeabilization buffer) for flow cytometry on ice. Intracellular staining was performed on the same or the next day using BD permeabilization buffer. See Table S1 for antibody probes used. For flow cytometry of leukocyte subsets, 3 million LNCs were taken immediately after harvest for extracellular staining and fixing. See Table S2 for antibody probes used. A BD Fortessa X20 was used for cellular phenotyping.

### Bone marrow-derived dendritic cells (BMDCs)

Bone marrow was extracted from mice by flushing femora and tibiae with PBS + 2% BSA + 2% glucose using a 25-G syringe. Red blood cells were lysed and remaining bone marrow cells were filtered through 70 µm cell strainer. Cells were cultured at 500,000 cells / ml R10 with 20 ng/ml murine rGM-CSF (PeproTech); cells were fed on day 3 with R10 + rGM-CSF, and on day 6 with R10 alone. On day 7, loosely adherent cells were harvested by gentle pipetting and used in assays. These cells were ∼60% CD11c+ MHC-II+ as we obtain routinely. To assess cytokine production, 100,000 BMDCs were transferred per well to 96-well plates precoated with GlcC14C18 (by dissolving in isopropanol and drying) and containing MDP in a final volume of 200 µl. BMDCs were incubated with stimulants for 6 hours at 37°C, 5% CO_2_ before supernatant was removed and stored at −80°C for later ELISA analysis (mouse TNF ELISA from ThermoScientific). To asses surface marker expression, 300,000 BMDCs were transferred per FACS tube precoated with 50 ng GlcC14C18 (by dissolving in isopropanol and drying) and containing 100 ng/ml MDP in a final volume of 200 µl. BMDCs were incubated with stimulants for 24 or 48 hours at 37°C, 5% CO_2_ before staining and fixing for flow cytometry analysis. See Table S3 for antibodies used in flow cytometry.

### Relapsing-remitting experimental autoimmune encephalomyelitis (RR-EAE)

Adjuvants were prepared similarly as above, but with myelin oligodendrocyte glycoprotein (MOG) used as antigen. Briefly, PBS solution containing 1 mg/ml MOG +/-600 μg/ml *N*-glycolyl MDP was emulsified with CFA (1 mg/ml *M. tuberculosis*) / IFA plus 200 μg/ml GlcC14C18 (or 20 μg/ml TDM) by back-and-forth extrusion through an 18 G two-way needle with two all-plastic syringes. 8-12 week old female C57BL/6 mice (Charles River) were induced for RR-EAE by a standard protocol: briefly, on day 0, pertussis toxin (PT) (200 ng) was administered i.v. in the tail vein, and mice were immunized with 50 μg MOG by bilateral s.c. injection in the back towards the tail with 100 μl total of an emulsification of CFA or IFA plus μ mycobacterial MAMPs (adjuvant group was randomized within and across cages). On day 2, mice received a second equivalent dose of PT i.v. After about one week, EAE was scored blinded (i.e. the scorer did not know the adjuvant received by the mouse) every day according to these cumulative criteria: 0, no paralysis (normal); 0.5, partial tail weakness observed as < 50% of tail dragging when mouse walks; 1, tail paralysis observed as >50% of tail dragging as mouse walks; 2, slow righting reflex (delay < 5 seconds) when mouse is flipped; 3, very slow or absent righting reflex (> 5 seconds) or inability to bear weight with back legs observed as dragging hindquarters when walking; 3.5, partial paralysis of one or both hind limbs; 4, complete paralysis of one or both hindlimbs; 4.5, complete paralysis of one or both hind limbs plus trunk weakness; 5, weakness or paralysis of forelimbs; 6, found dead. Mice were weighed every other day during scoring. Mice reaching a score of 5 were euthanized within 24 hours. Blinding was accomplished by having one person inject the mice and another person score/weigh the mice, unaware of experimental group.

### Histopathology

At the end of EAE scoring, the experiment was un-blinded and three mice from each group with matching scores were selected. These six mice were anesthetized with peritoneal ketamine injection, were perfused with PBS and formalin, and then spinal cords were extracted, equilibrated in sucrose and then frozen in OTC compound (VWR). Frozen tissue was sectioned 14-nm thick onto slides and then stained (Nissl or Luxol fast blue). Sections of good quality (20 per mouse, 5 cervical, 5 upper thoracic, 5 lower thoracic and 5 lumbar) were randomly selected and photographed with a Nikon Eclipse microscope. Randomized (blinded) Nissl stain photographs were used to quantify the area of disease in the spinal cord by subtracting the total 2D area of each tissue section with the area without cellular infiltration in white matter (thus, the difference of areas is the area with cellular infiltration). This was divided by the total area to determine the fraction or percent of diseased area.

### Software, data analysis and statistics

Flow cytometry data was acquired using FACSDiva^TM^ software (BD). FCS files were analyzed using FlowJo V10 (BD). Digital microscopy images were analyzed with ImageJ (NIH). Graphs were generated with, and routine statistical testing was accomplished with GraphPad Prism v7 or v8 (GraphPad Software Inc.). Assessment for normal data, then parametric or non-parametric analyses were applied, as indicated. Sample-size and power calculations were performed manually using Microsoft Excel. Manuscript figures were assembled with Microsoft PowerPoint.

## Acknowledgments

We would like to thank the Containment Level 3 platform of the Research Institute of the McGill University Health Centre (RI-MUHC) for providing the needed facilities and intellectual support. We thank the Immunophenotyping platform of the RI-MUHC for cytometers and advice. We are very grateful to Christine Wells (University of Melbourne, Australia) for providing *Mincle*-KO mice and to Alexiane Decout for preparing GlcC14C18. For tutelage and technical assistance, J.Y.D. is thankful to Damien Montamat-Sicotte, Ourania Tsatas, Sarah Danchuk, Iain Roe, Andréanne Lupien, Nimara Asbah, and Daniel Houle. J.Y.D. was personally supported by the Canadian Institutes of Health Research (CIHR) Canada Graduate Scholarship – Master’s Program, Fonds de Recherche du Québec – Santé (FRQ-S) Doctoral Training Award, RI-MUHC studentships (Master’s and Ph.D.) and scholarships from the McGill Department of Microbiology and Immunology. Research activities were funded by CIHR foundation grant held by M.B. and a CIHR grant held by S.D.

## Additional Information

Competing interests: the authors have none.

## Author Contributions

Conceptualization, J.Y.D. and M.B.; Methodology, J.Y.D. and J.G.Z.; Investigation, J.Y.D., F.M. and J.G.Z.; Resources, S.D., J.N. and M.B.; Writing – Original Draft, J.Y.D. and M.B.; Writing – Review & Editing, J.Y.D., F.M., J.G.Z., S.D., J.N. and M.B.; Visualization, J.Y.D.; Supervision, S.D. and M.B.; Funding acquisition, J.Y.D., S.D. and M.B.

## References

1. Chase, M. W. & Landsteiner, K. Experiments on Transfer of Cutaneous Sensitivity to Simple Compounds. Proc. Soc. Exp. Biol. Med. 49, 688–690 (1942).

2. Chase, M. W. The Cellular Transfer of Cutaneous Hypersensitivity to Tuberculin. Proc. Soc. Exp. Biol. Med. 59, 134–135 (1945).

3. Baxter, A. G. The origin and application of experimental autoimmune encephalomyelitis. Nat. Rev. Immunol. 7, 904–912 (2007).

4. Constantinescu, C. S., Farooqi, N., O’Brien, K. & Gran, B. Experimental autoimmune encephalomyelitis (EAE) as a model for multiple sclerosis (MS). Br. J. Pharmacol. 164, 1079–1106, doi:10.1111/bph.2011.164.issue-4 (2011).

5. Vogel, F. R. Improving Vaccine Performance with Adjuvants. Clin. Infect. Dis. 30, S266–270 (2000).

6. Adam, A., Ellouz, F., Ciorbaru, R., Petit, J. F. & Lederer, E. Peptidoglycan adjuvants: minimal structure required for activity. Z. Immunitatsforsch. Exp. Klin. Immunol. 149, 341–348 (1975).

7. Coulombe, F. et al. Increased NOD2-mediated recognition of *N*-glycolyl muramyl dipeptide. J. Exp. Med. 206, 1709–1716, doi:10.1084/jem.20081779 (2009).

8. Raymond, J. B., Mahapatra, S., Crick, D. C. & Pavelka, M. S., Jr. Identification of the namH gene, encoding the hydroxylase responsible for the N-glycolylation of the mycobacterial peptidoglycan. J. Biol. Chem. 280, 326–333, doi:10.1074/jbc.M411006200 (2005).

9. Mahapatra, S., Crick, D. C., McNeil, M. R. & Brennan, P. J. Unique structural features of the peptidoglycan of Mycobacterium leprae. J. Bacteriol. 190, 655–661, doi:10.1128/JB.00982-07 (2008).

10. Essers, L. & Schoop, H. J. Evidence for the incorporation of molecular oxygen, a pathway in biosynthesis of *N*-glycolylmuramic acid in *Mycobacterium phlei*. Biochim. Biophys. Acta 544, 180–184 (1978).

11. Hansen, J. M. et al. *N*-glycolylated peptidoglycan contributes to the immunogenicity but not pathogenicity of Mycobacterium tuberculosis. J. Infect. Dis. 209, 1045–1054, doi:10.1093/infdis/jit622 (2014).

12. Girardin, S. E. et al. Nod2 is a general sensor of peptidoglycan through muramyl dipeptide (MDP) detection. J. Biol. Chem. 278, 8869–8872, doi:10.1074/jbc.C200651200 (2003).

13. Miceli-Richard, C. et al. CARD15 mutations in Blau syndrome. Nat. Genet. 29, 19–20, doi:10.1038/ng720 (2001).

14. Ogura, Y. et al. A frameshiftmutation in NOD2 associated with susceptibility to Crohn’s disease. Nat. Lett. 411, 603–606 (2001).

15. Austin, C. M., Ma, X. & Graviss, E. A. Common nonsynonymous polymorphisms in the NOD2 gene are associated with resistance or susceptibility to tuberculosis disease in African Americans. J. Infect. Dis. 197, 1713–1716, doi:10.1086/588384 (2008).

16. Zhang, F. R. et al. Genomewide association study of leprosy. N. Engl. J. Med. 361, 2609–2618, doi:10.1056/NEJMoa0903753 (2009).

17. Shenderov, K. et al. Cord factor and peptidoglycan recapitulate the Th17-promoting adjuvant activity of mycobacteria through Mincle/CARD9 signaling and the inflammasome. J. Immunol. 190, 5722–5730, doi:10.4049/jimmunol.1203343 (2013).

18. Ishikawa, E. et al. Direct recognition of the mycobacterial glycolipid, trehalose dimycolate, by C-type lectin Mincle. J. Exp. Med. 206, 2879–2888, doi:10.1084/jem.20091750 (2009).

19. Miyake, Y. et al. C-type lectin MCL is an FcRgamma-coupled receptor that mediates the adjuvanticity of mycobacterial cord factor. Immunity 38, 1050–1062, doi:10.1016/j.immuni.2013.03.010 (2013).

20. Bekierkunst, A. Acute Granulomatous Response Produced in Mice by Trehalose-6,6-Dimycolate. J. Bacteriol. 96, 958–961 (1968).

21. Shigeru, K. et al. Synthesis of Muramyl Dipeptide Analogs with Enhanced Adjuvant Activity. Bull. Chem. Soc. Japan 53, 2570–2577 (1980).

22. Xing, S. & Gleason, J. L. A robust synthesis of N-glycolyl muramyl dipeptide via azidonitration/reduction. Org. Biomol. Chem. 13, 1515–1520, doi:10.1039/c4ob02147a (2015).

23. Decout, A. et al. Rational design of adjuvants targeting the C-type lectin Mincle. Proc. Natl. Acad. Sci. U. S. A. 114, 2675–2680, doi:10.1073/pnas.1612421114 (2017).

24. Divangahi, M. et al. NOD2-Deficient Mice Have Impaired Resistance to Mycobacterium tuberculosis Infection through Defective Innate and Adaptive Immunity. J. Immunol. 181, 7157–7165, doi:10.4049/jimmunol.181.10.7157 (2008).

25. Wells, C. A. et al. The Macrophage-Inducible C-Type Lectin, Mincle, Is an Essential Component of the Innate Immune Response to Candida albicans. The Journal of Immunology 180, 7404–7413, doi:10.4049/jimmunol.180.11.7404 (2008).

26. Merad, M., Sathe, P., Helft, J., Miller, J. & Mortha, A. The dendritic cell lineage: ontogeny and function of dendritic cells and their subsets in the steady state and the inflamed setting. Annu. Rev. Immunol. 31, 563–604, doi:10.1146/annurev-immunol-020711-074950 (2013).

27. Janeway, C. A. Approaching the Asymptote? Evolution and Revolution in Immunology. Cold Spring Harb. Symp. Quant. Biol. 54, 1–13 (1989).

28. Odendall, C. & Kagan, J. C. Host-Encoded Sensors of Bacteria: Our Windows into the Microbial World. MicrobiolSpectrum 7 (2019).

29. Yonekawa, A. et al. Dectin-2 Is a Direct Receptor for Mannose-Capped Lipoarabinomannan of Mycobacteria. Immunity 41, 402–413 (2014).

30. Decout, A. et al. Deciphering the molecular basis of mycobacteria and lipoglycan recognition by the C-type lectin Dectin-2. Sci. Rep. 8, 16840, doi:10.1038/s41598-018-35393-5 (2018).

31. Stamm, C. E., Collins, A. C. & Shiloh, M. U. Sensing of Mycobacterium tuberculosis and consequences to both host and bacillus. Immunol. Rev. 264, 204–219, doi:10.1111/imr.12263 (2015).

32. Prinz, M. et al. Innate immunity mediated by TLR9 modulates pathogenicity in an animal model of multiple sclerosis. J. Clin. Invest. 116, 456–464, doi:10.1172/JCI26078 (2006).

33. Reynolds, J. M. et al. Toll-like receptor 2 signaling in CD4(+) T lymphocytes promotes T helper 17 responses and regulates the pathogenesis of autoimmune disease. Immunity 32, 692–702, doi:10.1016/j.immuni.2010.04.010 (2010).

34. Miranda-Hernandez, S. et al. Role for MyD88, TLR2 and TLR9 but not TLR1, TLR4 or TLR6 in experimental autoimmune encephalomyelitis. J. Immunol. 187, 791–804, doi:10.4049/jimmunol.1001992 (2011).

35. Shaw, P. J. et al. Signaling via the RIP2 adaptor protein in central nervous system-infiltrating dendritic cells promotes inflammation and autoimmunity. Immunity 34, 75–84, doi:10.1016/j.immuni.2010.12.015 (2011).

36. Collins, Angela C. et al. Cyclic GMP-AMP Synthase Is an Innate Immune DNA Sensor for Mycobacterium tuberculosis. Cell Host & Microbe 17, 820–828, doi:10.1016/j.chom.2015.05.005 (2015).

37. Watson, Robert O. et al. The Cytosolic Sensor cGAS Detects Mycobacterium tuberculosis DNA to Induce Type I Interferons and Activate Autophagy. Cell Host & Microbe 17, 811–819, doi:10.1016/j.chom.2015.05.004 (2015).

38. Bafica, A. et al. TLR9 regulates Th1 responses and cooperates with TLR2 in mediating optimal resistance to Mycobacterium tuberculosis. J. Exp. Med. 202, 1715–1724, doi:10.1084/jem.20051782 (2005).

39. Chen, Z. et al. Association between toll-like receptors 9 (TLR9) gene polymorphism and risk of pulmonary tuberculosis: meta-analysis. BMC Pulm. Med. 15, 57, doi:10.1186/s12890-015-0049-4 (2015).

40. Eisenbarth, S. C. et al. Lipopolysaccharide-enhanced, toll-like receptor 4-dependent T helper cell type 2 responses to inhaled antigen. J. Exp. Med. 196, 1645–1651, doi:10.1084/jem.20021340 (2002).

41. Kobayashi, K. S. et al. Nod2-Dependent Regulation of Innate and Adaptive Immunity in the Intestinal Tract. Science 307, 731–734 (2004).

42. Khan, N. et al. Signaling through NOD-2 and TLR-4 Bolsters the T cell Priming Capability of Dendritic cells by Inducing Autophagy. Sci. Rep. 6, 19084, doi:10.1038/srep19084 (2016).

43. Wolfert, M. A., Murray, T. F., Boons, G. J. & Moore, J. N. The origin of the synergistic effect of muramyl dipeptide with endotoxin and peptidoglycan. J. Biol. Chem. 277, 39179–39186, doi:10.1074/jbc.M204885200 (2002).

44. Allison, D. G. & Lambert, P.A. in Molecular Medical Microbiology Ch. 32, 583–598 (2015).

45. Berard, J. L., Wolak, K., Fournier, S. & David, S. Characterization of relapsing-remitting and chronic forms of experimental autoimmune encephalomyelitis in C57BL/6 mice. Glia 58, 434–445, doi:10.1002/glia.20935 (2010).

46. Brand, D. D., Latham, K. A. & Rosloniec, E. F. Collagen-induced arthritis. Nat. Protoc. 2, 1269–1275, doi:10.1038/nprot.2007.173 (2007).

47. Kleinnijenhuis, J. et al. Bacille Calmette-Guerin induces NOD2-dependent nonspecific protection from reinfection via epigenetic reprogramming of monocytes. Proc. Natl. Acad. Sci. U. S. A. 109, 17537–17542, doi:10.1073/pnas.1202870109 (2012).

48. Wong, S. Y. et al. B Cell Defects Observed in Nod2 Knockout Mice Are a Consequence of a Dock2 Mutation Frequently Found in Inbred Strains. J. Immunol. 201, 1442–1451, doi:10.4049/jimmunol.1800014 (2018).

