## Supplemental text and figures for "Synthetic mycobacterial molecular patterns partially complete Freund’s adjuvant"

Supplemental Figure Titles and Legends

**Figure S1. Flow cytometry gating strategies for lymph node cells. A,** gating to quantify cytokine-producing CD4+CD8- T cells. Shown are representative gating and data from an OVA-stimulated sample (CFA-immunized WT mouse). For comparison, cytokine production for the corresponding unstimulated sample is included. **B**, gating to identify lymph node cell subsets. Shown is gating on a representative sample (CFA-immunized WT mouse). pDCs are B220+Ly6C+MHC-II+CD11b-CD11c+ cells. B cells are B220+Ly6C-CD11b-CD11c-CD4-CD8a- cells. cDCs are B220-MHC-II$^{hi}$Ly6C-CD11c+ cells (analyzed subsets are CD11b+/-). CD4+ T cells are B220-CD11b-CD4+CD8a- cells. CD8+ T cells are B220-CD11b-CD4-CD8a+ cells. Monocytes are B220-CD11b+Ly6C+SSC$^{lo}$ cells. PMNs are B220-CD11b+Ly6C$^{med}$SSC$^{hi}$ CD11c- cells.

**Figure S2. Addition data on CFA-dependent cell-mediated immune responses as a function of mycobacterial *namH*.** These data are from the same experiments shown in fig. 1C (refer to fig. 1C legend for details). **A-B**, Total numbers (from two lymph nodes) of cytokine-producing CD4+CD8- lymph node cells of mice immunized against OVA with H37Rv, H37Rv Δ*namH*, or IFA alone. Shown are averages +/- SEM. p-values were calculated with two-tailed student's *t*-tests. **p<0.01. **C-D**, Contribution of *namH* to the mycobacterial portion of OVA-specific IFN-γ elicited by CFA. **C**, result was obtained from %IFN-γ+ data by subtracting the average IFA background from all CFA data, and plotting the results as % of IFA+H37Rv 'wild-type'. **D**, result was obtained from # IFN-γ+ data by subtracting the average IFA background from all CFA data, and plotting the results as % of IFA+H37Rv 'wild-type'. Shown are averages +/- SEM. p-values were calculated with two-tailed student's t-tests. *p<0.05. **E**, results separated by experimental run for the data presented in fig. 1C and fig. S2A-B. Shown are averages +/- SEM.

**Figure S3. CFA-dependent IFN-γ response by ELISpot as a function of host *Nod2*. A-B**, IFN-γ ELISpot of inguinal lymph node cells mice immunized against OVA seven days prior, produced in an independent experiment. **A**, number of IFN-γ spot-forming cells per one million cells. **B**, total number of IFN-γ spot-forming cells per two inguinal lymph nodes. Shown are averages +/- SEM. p-values were calculated with two-tailed student's *t*-tests. *p<0.05. For CFA *Nod2*+/+, CFA *Nod2*-/-, IFA *Nod2*+/+ and IFA *Nod2*-/-, N = 6, 8, 6 and 6 mice, respectively. **C**, flow cytometry of inguinal lymph node cells mice immunized against OVA seven days prior, produced in an independent experiment similar to that in fig. 2A-B. **D**, results separated by experimental run for the data presented in fig. 2C-D and fig. S4C-D. Shown are averages +/- SEM.

**Figure S4. Additional data on CFA-dependent cell-mediated immune responses as a function of host *Nod2* and Mincle.** These data are from the same experiments shown in fig. 2

(refer to fig. 2 legend for details). **A-F**, Total numbers (from two lymph nodes) of cytokine-producing CD4+CD8- lymph node cells of mice immunized against OVA with the indicated adjuvant as a function of: **A-B**, *Nod2*; **C-D**, *Mincle*; **E-F**, *Mincle* and *Nod2* together. Shown are averages +/- SEM. p-values were calculated with two-tailed student's *t*-tests. *$p<0.05$; ns, not significant, $p>0.05$.

**Figure S5. Additional data on MHC-II expression and costimulatory molecule upregulation by BMDCs stimulated with GlcC14C18 and MDPs**. **A**, percentage of cells expressing MHC-II at high levels (according to gate in fig. 3B) after 48 hours of stimulation with the indicated MAMPs. **B-E**, median fluorescence intensity of **B**, MHC-II; **C**, CD40; **D**, CD80; **E**, CD86 on CD11b+CD11c+MHC-II$^{hi}$ cells after 48 hours of stimulation with the indicated MAMPs (use legend of panel A). Shown are averages +/- SD of 3 individually stimulated and assayed cultures. **F-I**, histograms of CD11b+CD11c+MHC-II$^{hi}$ cells demonstrating expression levels of **F**, MHC-II; **G**, CD40; **H**, CD80; **I**, CD86, for both timepoints (FMOC is not shown for MHC-II because the analysis required MHC-II gating).

**Figure S6. Synthetic adjuvant-dependent IFN-γ responses compared to CFA by ELISpot.** **A-B**, IFN-γ ELISpot of inguinal lymph node cells of mice immunized against OVA with the indicated adjuvant seven days prior. **A**, number of IFN-γ spot-forming cells per one million cells. **B**, total number of IFN-γ spot-forming cells per two inguinal lymph nodes. Data was pooled from two independent experiments. Shown are averages +/- SEM, where for IFA, IFA + 3 µg *N*-glycolyl MDP, IFA + 10 µg *N*-glycolyl MDP, IFA + 30 µg *N*-glycolyl MDP, IFA + 100 µg *N*-glycolyl MDP and CFA, N = 11, 6, 12, 11, 6 and 11 mice, respectively. **C-D**, These data are from the same mice used in fig. 4B (refer to fig. 4B legend for details). **C**, number of IFN-γ spot-forming cells per one million cells. **D**, total number of IFN-γ spot-forming cells per two inguinal lymph nodes. Shown are averages +/- SEM. In comparing IFA + 30 µg MDP to IFA + MDP + 10 or 30 µg GlcC14C18, p-values were calculated using Dunnett's T3 multiple comparisons test. *$p<0.05$; **$p<0.01$; ns, not significant, $p>0.05$.

**Figure S7. Lymph node cell subset numbers after immunization with synthetic adjuvants.** These data are from the same mice used in fig. 5 (refer to fig. 5 legend for details of the experiment). **A-F,** From two inguinal lymph nodes at 4 or 7 days post-immunization, shown are total numbers of extracted **A**, B cells; **B**, CD4+ T cells; **C**, CD8+ T cells; **D**, plasmacytoid dendritic cells (pDCs); **E**, monocytes; **F**, polymorphonuclear cells (PMNs). Data are graphed as averages +/- SEM.

**Figure S8. Additional statistics for RR-EAE.** These data are from the same set shown in fig. 5 (refer to fig. 5 legend for details). **A**, average weight of mice over time +/- SEM. **B**, maximum EAE score reached on any day by mice, as of day 28. **C**, disease course of mice selected for spinal cord histopathology. **D-G**, Statistics for RR-EAE induced by IFA+TDM+MDP. **D**, average EAE score +/- SEM over time of mice induced with CFA (N=13) or IFA + 1 µg TDM + 30 µg *N*-glycolyl MDP (N=13). Mice were euthanized on day 27 post injection. **E**, cumulative EAE score, obtained by adding the EAE score of each mouse over each of the 27 days. Lines represent averages +/- SEM. **F**, average weight of mice over time +/- SEM. **G**, maximum EAE score reached on any day by mice, as of day 27.

Supplemental Tables

**Table S1. Flow cytometry antibodies for lymph node cell cytokines**

| Target | Supplier | Clone | Fluorochrome* |
|---|---|---|---|
| CD3ε | BD | 145-2C11 | PE |
| CD4 | BD | GK1.5 | BV786 |
| CD8α | BD | 53-6.7 | BV711 |
| CD19 | Biolegend | 6D5 | PE-Dazzle594 |
| B220/CD45R | BD | RA3-6B2 | BUV737 |
| IFN-γ | Biolegend | XMG1.2 | APC |
| IL-2 | BD | JES6-5H4 | BV605 |
| IL-4 | eBiosciences | BVD6-24G2 | PE-Cy7 |
| IL-17A | BD | TC11-18H10 | BUV395 |
| IL-10 | BD | JES5-16E3 | FITC |

*Viability dye: LIVE/DEAD™ Fixable Violet Dead Cell Stain (ThermoFisher Scientific)

**Table S2. Flow cytometry antibodies for lymph node cell DC and subset analysis**

| Target | Supplier | Clone | Fluorochrome* |
|---|---|---|---|
| B220/CD45R | BD | RA3-6B2 | BUV737 |
| CD4 | BD | GK1.5 | BV786 |
| CD8α | BD | 53-6.7 | BV711 |
| CD11b | Biolegend | M1/70 | BV605 |
| CD11c | BD | HL3 | FITC |
| CD40 | Biolegend | 3/23 | PE-Dazzle594 |
| CD80 | Biolegend | 16-10A1 | PE |
| CD86 | Biolegend | GL1 | PE-Cy7 |
| CD209 | BD | 5H10 | BUV395 |
| F4/80 | Biolegend | BM8 | APC |
| Ly6C | Biolegend | HK1.4 | APC-Cy7 |
| MHC-I (H-2$K^b$) | Biolegend | AF6-88.5 | BV510 |
| MHC-II (I-$A^b$) | Biolegend | AF6-120.1 | PerCP-Cy5.5 |

*Viability dye: LIVE/DEAD™ Fixable Violet Dead Cell Stain (ThermoFisher Scientific)

**Table S3. Flow cytometry antibodies for BMDCs**

| Target | Supplier | Clone | Fluorochrome* |
|---|---|---|---|
| CD11b | Biolegend | M1/70 | PerCP-Cy5.5 |
| CD11c | BD | HL3 | FITC |
| CD40 | Biolegend | 3/23 | PE-Dazzle594 |
| CD80 | Biolegend | 16-10A1 | PE |
| CD86 | Biolegend | GL1 | PE-Cy7 |
| MHC-II (I-$A^b$) | Biolegend | AF6-120.1 | APC |

*Viability dye: LIVE/DEAD™ Fixable Violet Dead Cell Stain (ThermoFisher Scientific)

### FIG. S1

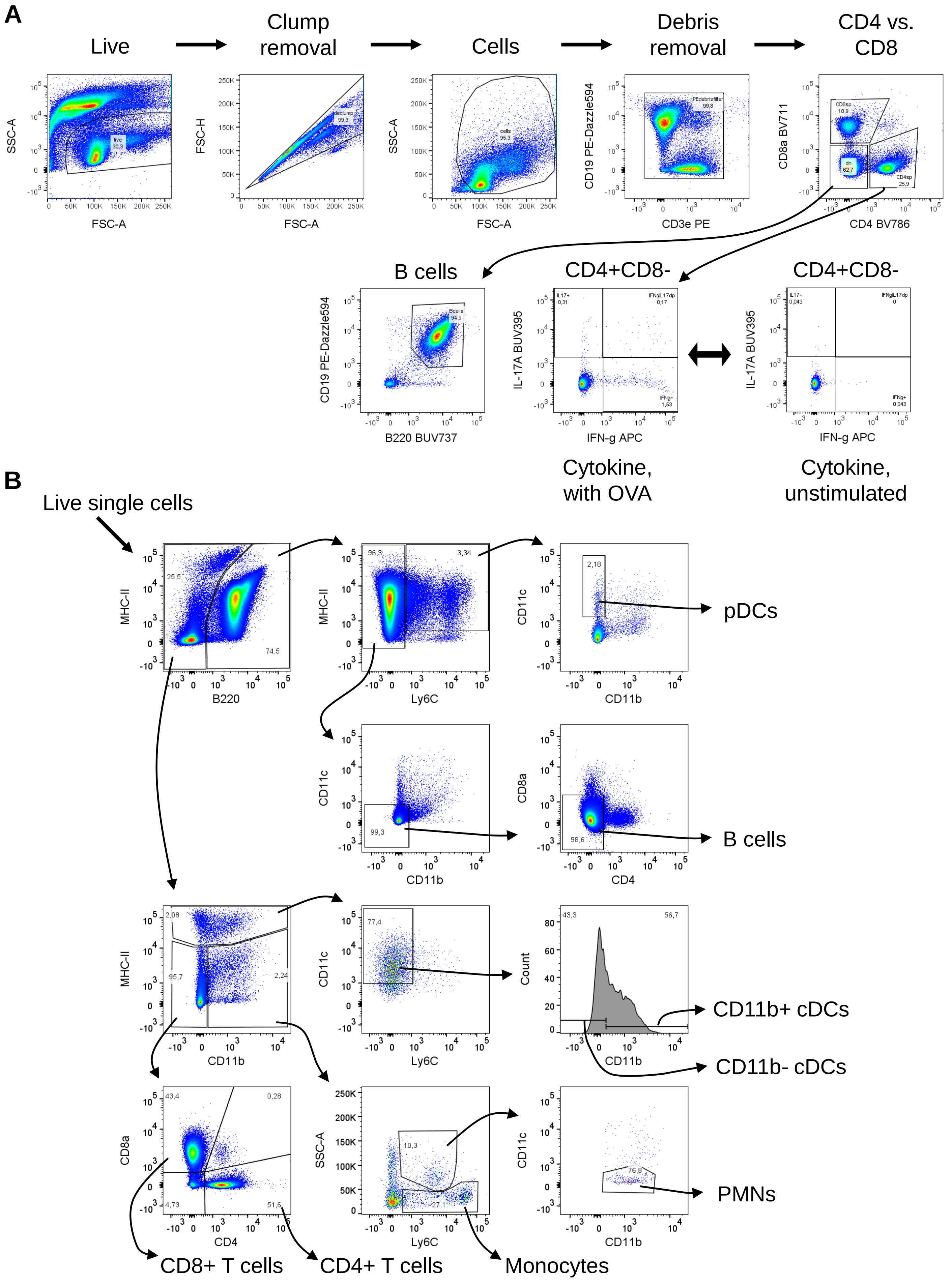

A
Live
Clump removal
Cells
Debris removal
CD4 vs. CD8
SSC-A
FSC-A
live 30,3
FSC-H
declump 99,3
cells 95,3
CD19 PE-Dazzle594
CD3e PE
PEdebrisfilter 99,8
CD8a BV711
CD4 BV786
CD8sp 10,9
dn 62,7
CD4sp 25,9
B cells
Bcells 94,9
B220 BUV737
CD4+CD8-
IL-17A BUV395
IFN-g APC
IL17+ 0,31
IFNgIL17dp 0,17
IFNg+ 1,53
IL17+ 0,043
IFNgIL17dp 0
IFNg+ 0,043
Cytokine, with OVA
Cytokine, unstimulated
B
Live single cells
MHC-II
B220
25,5
74,5
Ly6C
96,3
3,34
CD11c
CD11b
2,18
pDCs
99,3
CD8a
CD4
98,6
B cells
2,08
95,7
2,24
77,4
Count
43,3
56,7
CD11b+ cDCs
CD11b- cDCs
43,4
0,28
4,73
51,6
SSC-A
10,3
27,1
76,8
PMNs
CD8+ T cells
CD4+ T cells
Monocytes

### FIG. S2

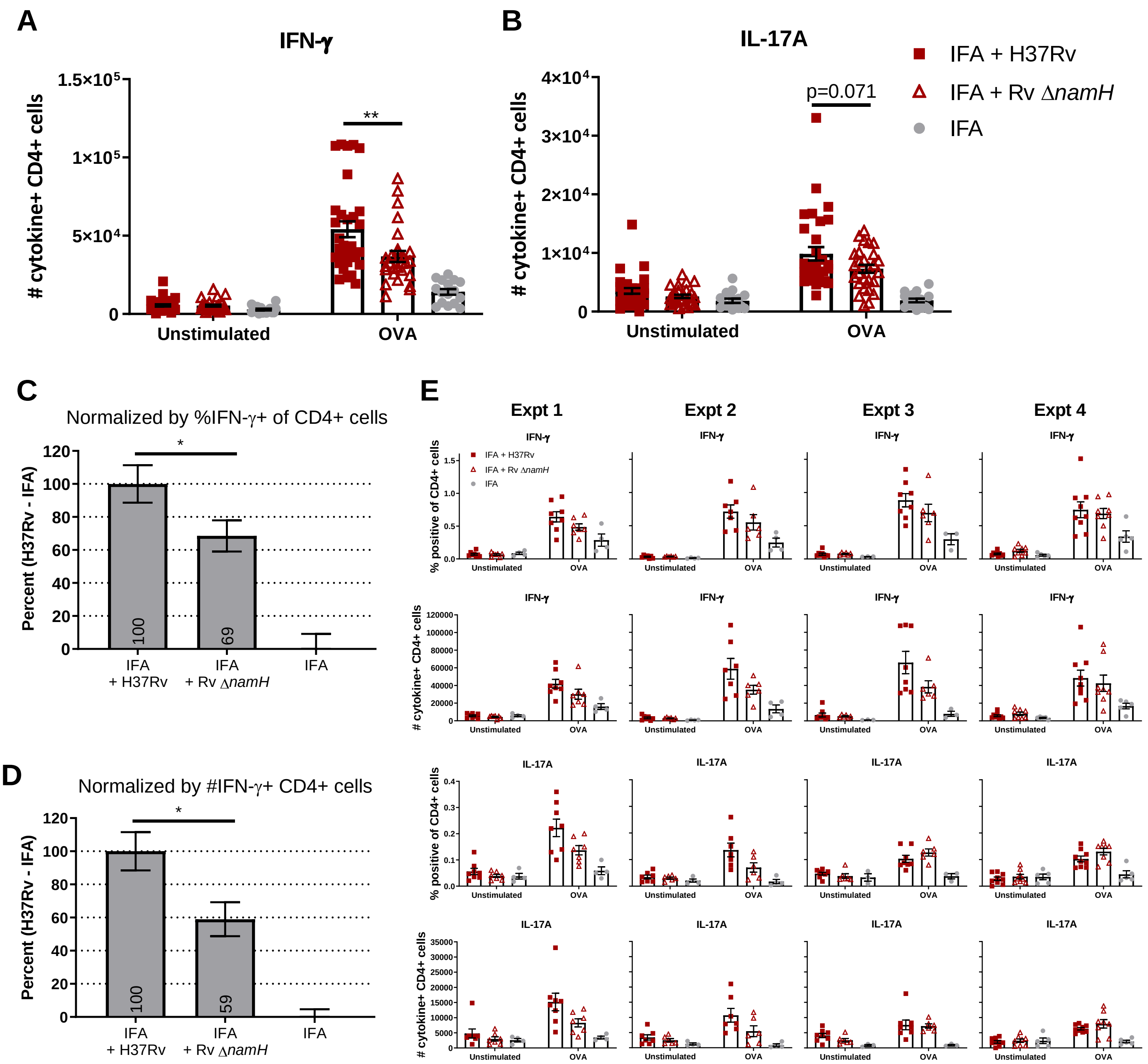

A
IFN-γ
### cytokine+ CD4+ cells
1.5×10^5
1×10^5
5×10^4
0
**
Unstimulated
OVA
B
IL-17A
4×10^4
3×10^4
2×10^4
1×10^4
0
p=0.071
IFA + H37Rv
IFA + Rv ΔnamH
IFA
C
Normalized by %IFN-γ+ of CD4+ cells
Percent (H37Rv - IFA)
120
100
80
60
40
20
0
*
100
69
IFA + H37Rv
IFA + Rv ΔnamH
IFA
D
Normalized by #IFN-γ+ CD4+ cells
*
100
59
E
Expt 1
Expt 2
Expt 3
Expt 4
IFN-γ
IL-17A
% positive of CD4+ cells
### cytokine+ CD4+ cells

FIG. S3

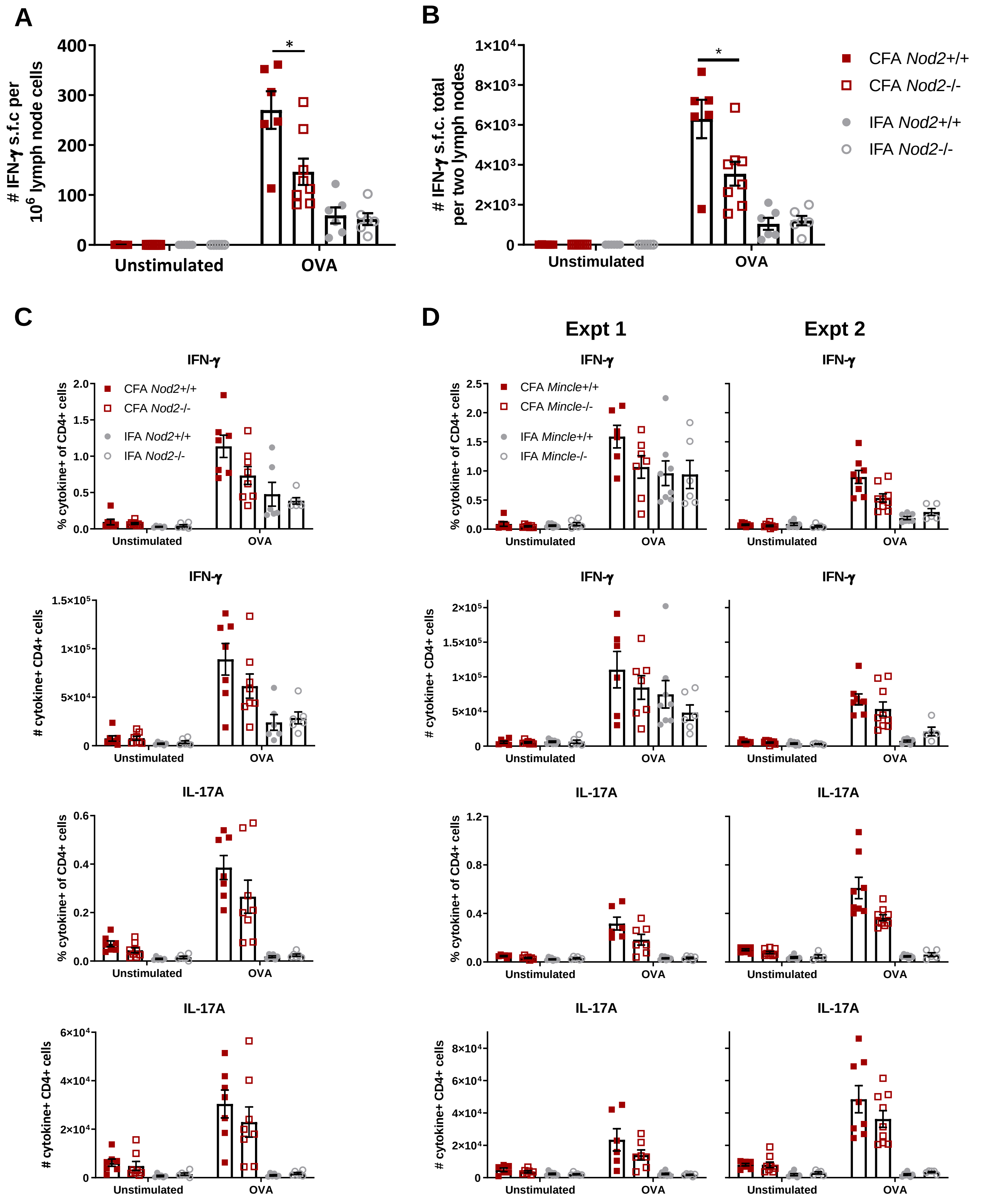

A
### IFN-γ s.f.c per $10^6$ lymph node cells
400
300
200
100
0
*
Unstimulated
OVA
B
### IFN-γ s.f.c. total per two lymph nodes
1×10⁴
8×10³
6×10³
4×10³
2×10³
0
*
Unstimulated
OVA
CFA Nod2+/+
CFA Nod2-/-
IFA Nod2+/+
IFA Nod2-/-
C
IFN-γ
% cytokine+ of CD4+ cells
CFA Nod2+/+
CFA Nod2-/-
IFA Nod2+/+
IFA Nod2-/-
Unstimulated
OVA
IFN-γ
### cytokine+ CD4+ cells
IL-17A
% cytokine+ of CD4+ cells
IL-17A
### cytokine+ CD4+ cells
D
Expt 1
Expt 2
IFN-γ
% cytokine+ of CD4+ cells
CFA Mincle+/+
CFA Mincle-/-
IFA Mincle+/+
IFA Mincle-/-
Unstimulated
OVA
IFN-γ
### cytokine+ CD4+ cells
IL-17A
% cytokine+ of CD4+ cells
IL-17A
### cytokine+ CD4+ cells

### FIG. S4

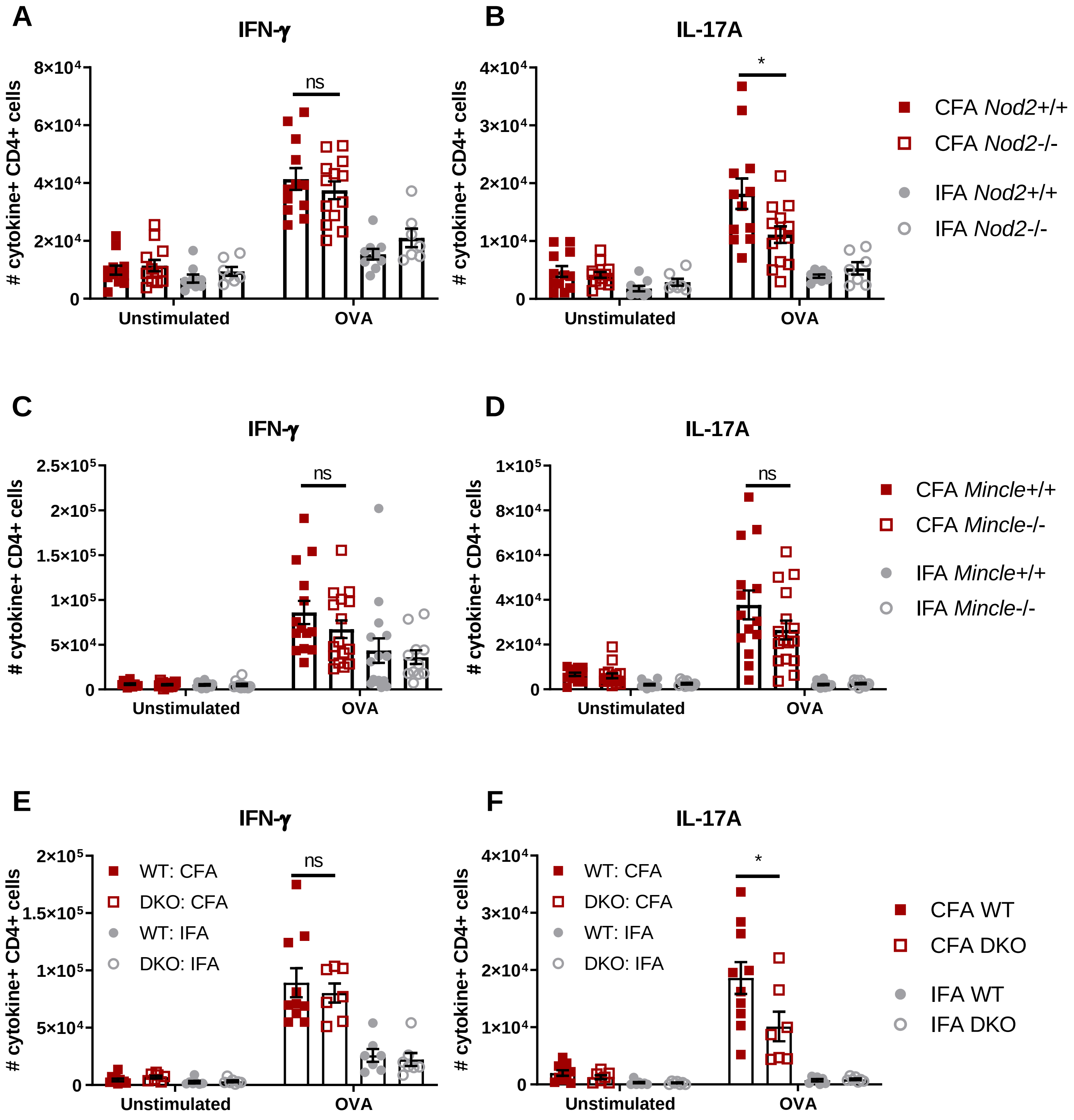

A
IFN-γ
### cytokine+ CD4+ cells
8×10⁴
6×10⁴
4×10⁴
2×10⁴
0
ns
Unstimulated
OVA
B
IL-17A
### cytokine+ CD4+ cells
4×10⁴
3×10⁴
2×10⁴
1×10⁴
0
*
Unstimulated
OVA
CFA Nod2+/+
CFA Nod2-/-
IFA Nod2+/+
IFA Nod2-/-
C
IFN-γ
### cytokine+ CD4+ cells
2.5×10⁵
2×10⁵
1.5×10⁵
1×10⁵
5×10⁴
0
ns
Unstimulated
OVA
D
IL-17A
### cytokine+ CD4+ cells
1×10⁵
8×10⁴
6×10⁴
4×10⁴
2×10⁴
0
ns
Unstimulated
OVA
CFA Mincle+/+
CFA Mincle-/-
IFA Mincle+/+
IFA Mincle-/-
E
IFN-γ
### cytokine+ CD4+ cells
2×10⁵
1.5×10⁵
1×10⁵
5×10⁴
0
WT: CFA
DKO: CFA
WT: IFA
DKO: IFA
ns
Unstimulated
OVA
F
IL-17A
### cytokine+ CD4+ cells
4×10⁴
3×10⁴
2×10⁴
1×10⁴
0
WT: CFA
DKO: CFA
WT: IFA
DKO: IFA
*
Unstimulated
OVA
CFA WT
CFA DKO
IFA WT
IFA DKO

### FIG. S5

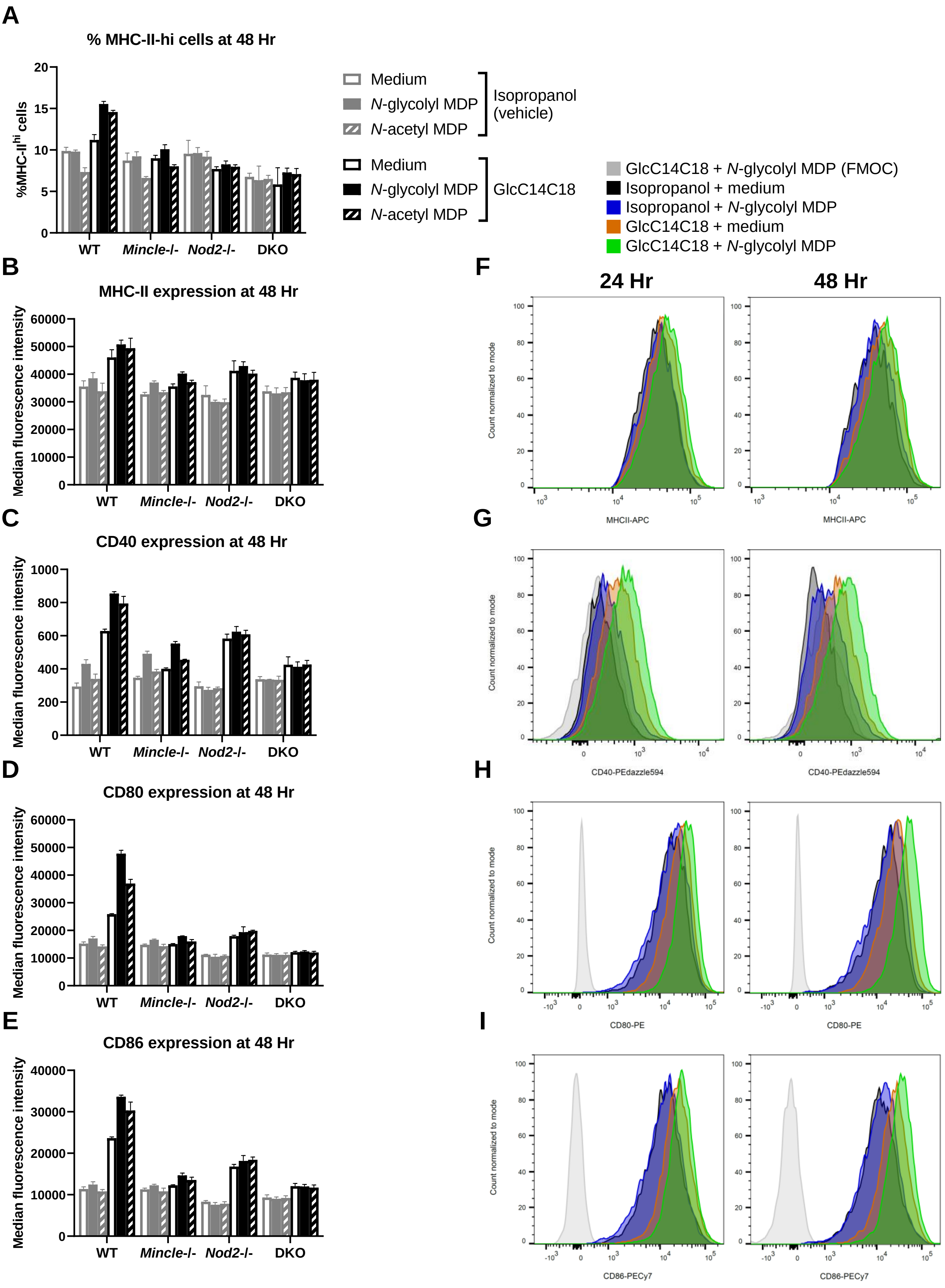
A
% MHC-II-hi cells at 48 Hr
%MHC-II^hi cells
WT
Mincle-/-
Nod2-/-
DKO
Medium
N-glycolyl MDP
N-acetyl MDP
Isopropanol (vehicle)
GlcC14C18
GlcC14C18 + N-glycolyl MDP (FMOC)
Isopropanol + medium
Isopropanol + N-glycolyl MDP
GlcC14C18 + medium
GlcC14C18 + N-glycolyl MDP
B
MHC-II expression at 48 Hr
Median fluorescence intensity
C
CD40 expression at 48 Hr
D
CD80 expression at 48 Hr
E
CD86 expression at 48 Hr
F
24 Hr
48 Hr
Count normalized to mode
MHCII-APC
G
CD40-PEdazzle594
H
CD80-PE
I
CD86-PECy7

### FIG. S6

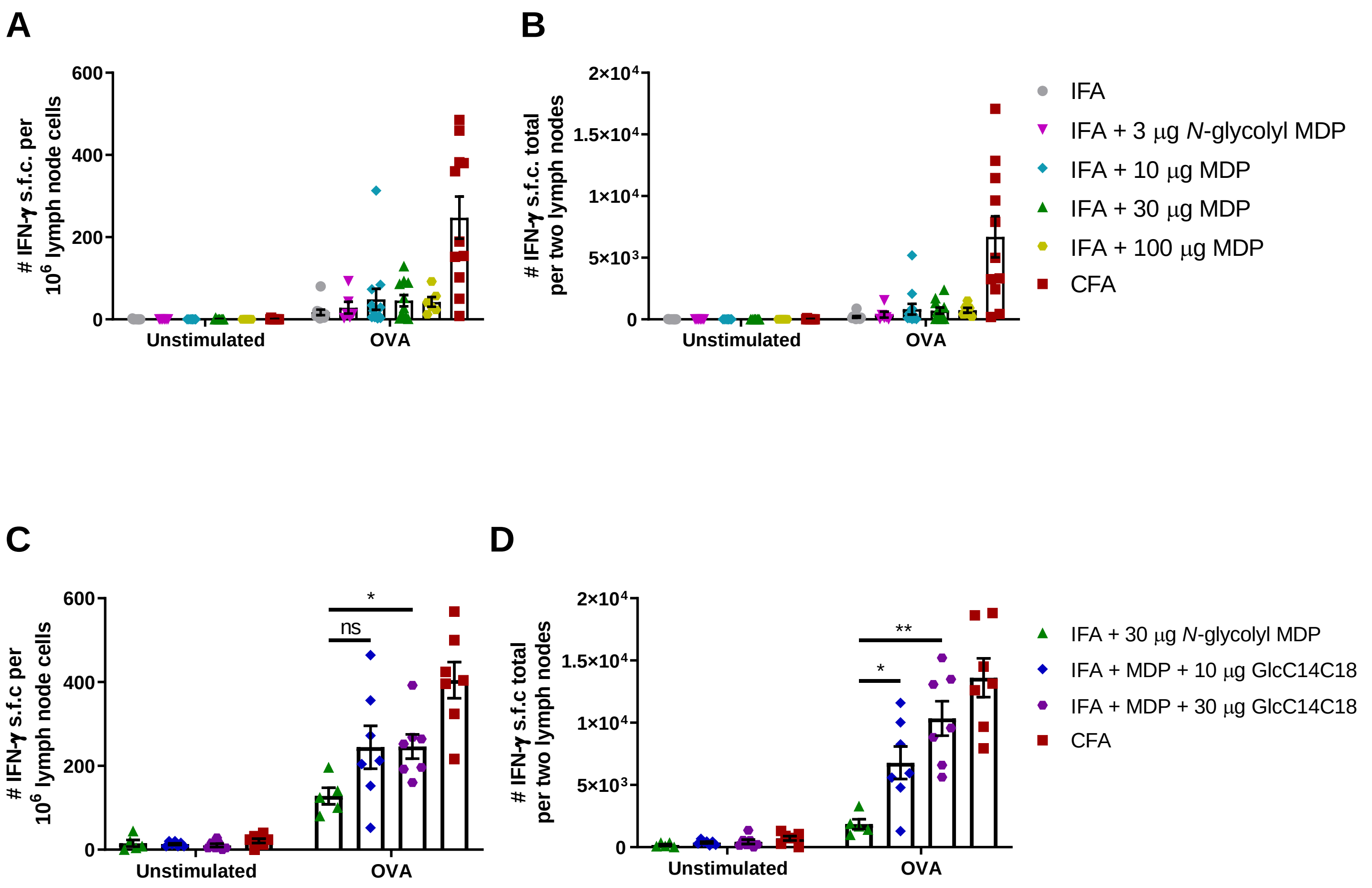
A
B
C
D
### IFN-γ s.f.c. per 10^6 lymph node cells
### IFN-γ s.f.c. total per two lymph nodes
### IFN-γ s.f.c per 10^6 lymph node cells
### IFN-γ s.f.c total per two lymph nodes
600
400
200
0
2×10^4
1.5×10^4
1×10^4
5×10^3
Unstimulated
OVA
ns
*
**
IFA
IFA + 3 μg N-glycolyl MDP
IFA + 10 μg MDP
IFA + 30 μg MDP
IFA + 100 μg MDP
CFA
IFA + 30 μg N-glycolyl MDP
IFA + MDP + 10 μg GlcC14C18
IFA + MDP + 30 μg GlcC14C18
CFA

### FIG. S7

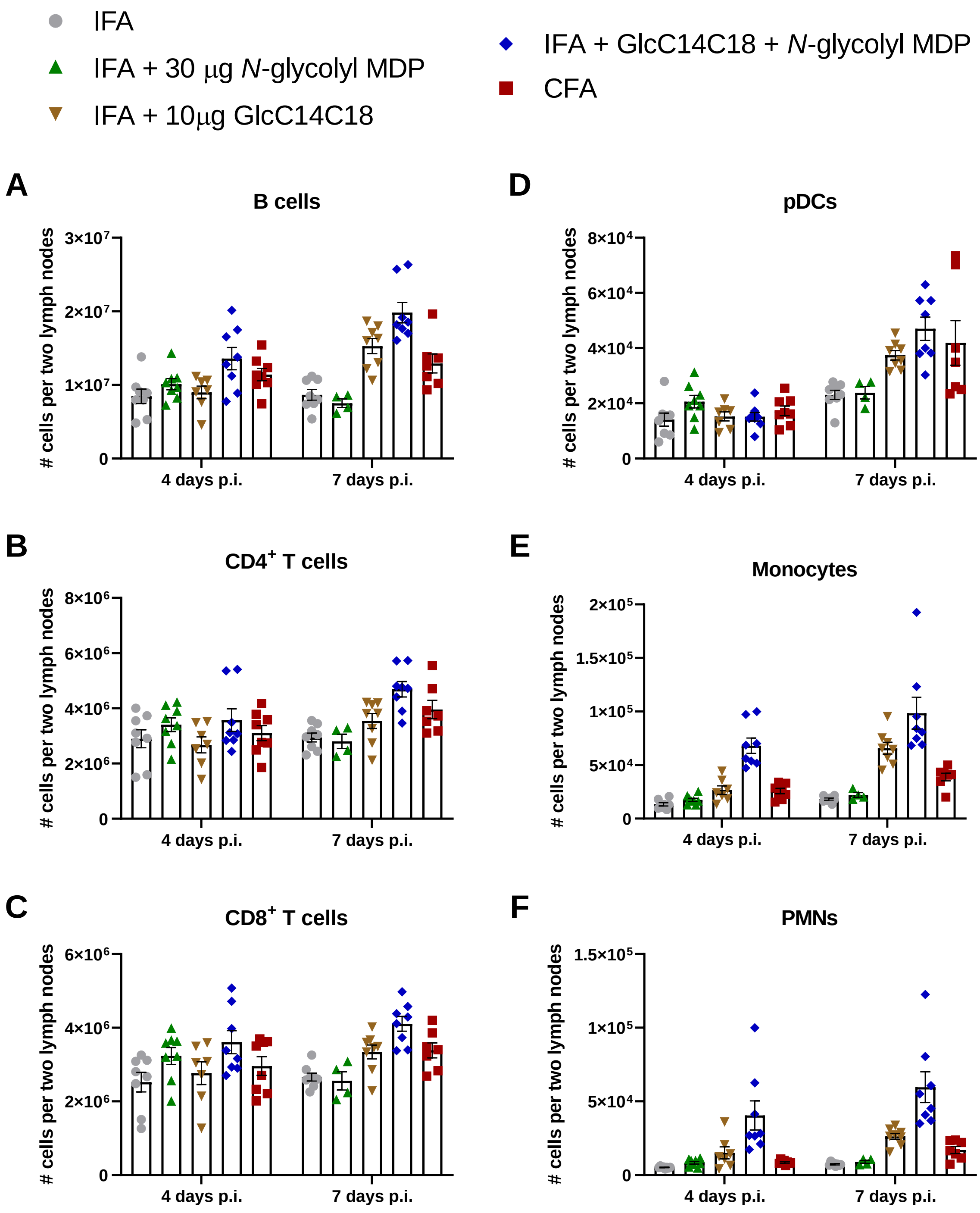

IFA
IFA + 30 μg N-glycolyl MDP
IFA + 10μg GlcC14C18
IFA + GlcC14C18 + N-glycolyl MDP
CFA
A
B cells
### cells per two lymph nodes
3×10^7
2×10^7
1×10^7
0
4 days p.i.
7 days p.i.
D
pDCs
8×10^4
6×10^4
4×10^4
2×10^4
B
CD4+ T cells
8×10^6
6×10^6
4×10^6
2×10^6
E
Monocytes
2×10^5
1.5×10^5
1×10^5
5×10^4
C
CD8+ T cells
6×10^6
4×10^6
2×10^6
F
PMNs
1.5×10^5
1×10^5
5×10^4

### FIG. S8

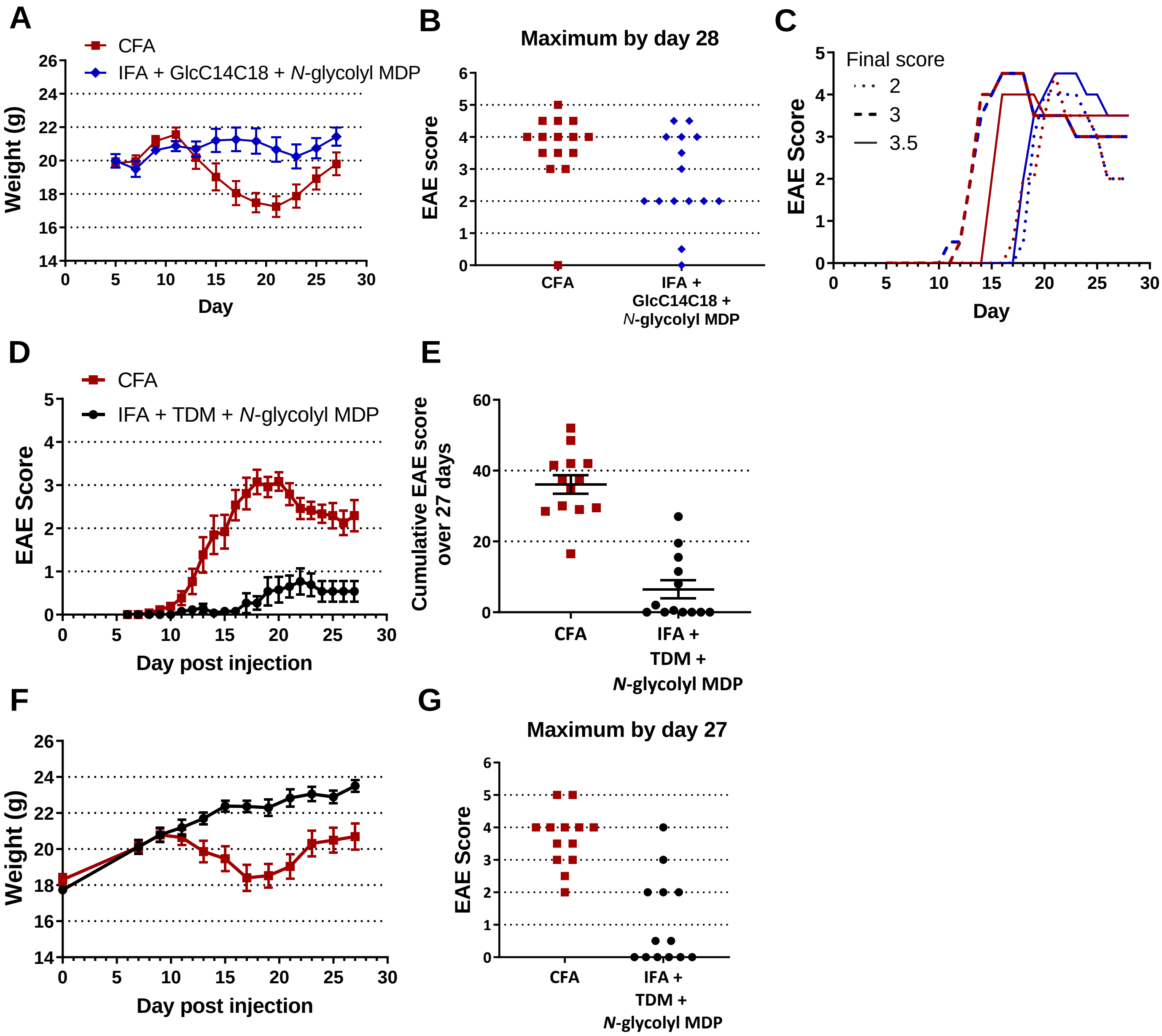
A
CFA
IFA + GlcC14C18 + N-glycolyl MDP
Weight (g)
Day
B
Maximum by day 28
EAE score
CFA
IFA + GlcC14C18 + N-glycolyl MDP
C
Final score
2
3
3.5
EAE Score
Day
D
CFA
IFA + TDM + N-glycolyl MDP
EAE Score
Day post injection
E
Cumulative EAE score over 27 days
CFA
IFA + TDM + N-glycolyl MDP
F
Weight (g)
Day post injection
G
Maximum by day 27
EAE Score
CFA
IFA + TDM + N-glycolyl MDP
